# Characterization of the knock-in *App*^SAA^ mouse model as a powerful tool for Alzheimer’s disease preclinical research

**DOI:** 10.64898/2026.09.08.750179

**Authors:** Lucy J. Sloan, Dylan T. Garceau, Nick J. Ryan, Justice Pomeroy-Tuck, Ritesh Chidambaram, Tim Ragan, Ravi S. Pandey, Erik B. Bloss, Gregory W. Carter, Adrian L. Oblak, Michael Sasner

## Abstract

**BACKGROUND:** Preclinical Alzheimer’s disease (AD) research requires models with age-dependent, physiologically relevant amyloid pathology, unlike overexpression in transgenic lines. We characterized the humanized *App*^SAA^ knock-in model to define translationally relevant therapeutic windows.

**METHODS:** Homozygous *App*^SAA^ and control mice were evaluated from 4 to 19 months using whole-brain 3D plaque imaging, cerebral amyloid angiopathy (CAA) quantification, multiplex neuropathology, brain transcriptomics, and plasma biomarkers. Hippocampal synaptic analysis and context-dependent memory was performed at 12 months.

**RESULTS:** Plaques, CAA, plaque-associated neuroinflammation, and dystrophic neurites accumulated with age. By 12 months, brain transcriptomes aligned with human AMP-AD co-expression modules, and plasma pTau-217, GFAP, and SNAP25 rose with age. *App*^SAA^ mice retained associative memory but showed memory interference and CA1 dendritic-spine loss that was exacerbated in proximity to plaques.

**CONCLUSIONS:** *App*^SAA^ mice recapitulate age-dependent, human-relevant AD phenotypes with defined therapeutic windows for preclinical research. The model is available without restriction from The Jackson Laboratory.

## 1. BACKGROUND

Alzheimer’s disease (AD) is a devastating and debilitating neurodegenerative disease, with no cure and available treatments providing only limited efficacy [1]. The development of efficacious clinical interventions has been hindered by the lack of animal models that reliably and appropriately recapitulate age-dependent disease progression [2]. In animal models, the therapeutic window for preclinical testing is minimized by transgenic overexpression of familial AD-associated genes such as amyloid precursor protein (APP) with familial AD (FAD)-associated mutations under cell-type specific artificial promoters [3–5], causing early over-expression of pathology. Additionally, transgenic models are plagued with confounding variables [2] such as highly variable and artifactually sex-specific disease pathology present in 5xFAD [6, 7] and APP/PS1 models [4, 8]. The development of knock-in animal models that more closely recapitulate age-dependent aspects of complex human disease processes are critical in bridging the translational research gap in the development of novel diagnostic biomarkers and interventional therapeutics [9, 10].

To address the gap between transgenic animal models and human clinical trials, the knock-in B6J.APP^SAA^ (*App*^SAA^) mouse model [11] was created. This model expresses a humanized Aβ fragment in the mouse *App* gene locus with the endogenous promoter along with the Swedish, Arctic, and Austrian mutations. Pathology develops in an age-dependent manner, despite significantly reduced levels of Aβ compared to the artificially elevated levels of amyloid expression in transgenic models. The *App*^SAA^ is commercially available with no restrictions on use, and characterization data presented here are publicly available to promote open science and serve as a reference for preclinical testing. This model aids in preclinical target identification, investigations into environmental and lifestyle risk factors, and serves as the genetic background for additional genetic risk factors such as tau and APOE, developing more complex pathologies to better understand human disease processes.

Importantly, this model is already being used for human-relevant preclinical research. Several groups evaluated the effects of anti-Aβ antibodies in the *App*^SAA^ model [12, 13] and non-amyloid therapeutic targets including Trem2 [14], septins [15], and erythropoietin [16]. Erythropoietin has been targeted in clinical trials internationally [17, 18], and results from the *App*^SAA^ mouse model are consistent with human data and results. Other preclinical studies have evaluated novel microglial imaging tracers [19, 20], and evaluated mechanistic roles of proteins like PHGDH [21] and CRMP2 [22]. Many of these studies replicate data presented previously [11], however several studies have reported novel phenotypes [23]. One study found that *App*^SAA^ mice not only recapitulate circadian instability and sleep dysregulation that is present in primarily female AD patients, but also showed that glial modulation improved non-cognitive deficits in female *App*^SAA^ mice [24].

To aid in preclinical development, the Model Organism Development and Evaluation for Late-Onset Alzheimer’s Disease (MODEL-AD) consortium has conducted an in-depth phenotyping evaluation of the *App*^SAA^ knock-in mouse model. While the original publication of *App*^SAA^ mice [11] established amyloid neuropathology and disease-associated microglia (DAM) phenotype, this work extends the phenotypic analysis to establish age-dependent disease-relevant therapeutic windows in line with research priorities [25]. Unlike transgenic models, the *App*^SAA^ model reliably and consistently accumulates amyloid plaques in an age-dependent manner. Brain transcriptomic data reflects an age-dependent increase in alignment between *App*^SAA^ mice and clinical data and clinically relevant plasma biomarkers showed age-dependent increases in markers of tau pathology, neuroinflammation and synaptic loss. Direct analysis of synapse loss was apparent at 12-months-of-age that was exacerbated in proximity to plaques and accompanied by memory deficits. Collectively, this data highlights the utility of this knock-in model over transgenic mouse models for preclinical research. Age-associated pathology allows for identification of therapeutic windows for various types of AD-associated pathology. Because disease mechanisms are age-dependent and age is the greatest risk factor for developing AD, this model is a fundamental advance, improving rigor and reproducibility of therapeutic development for the treatment of AD.

## 2. METHODS

### 2.1. Animals

All experiments were approved by the Institutional Animal Care and Use Committee (IACUC) at The Jackson Laboratory (IACUC #18051 and #20006). Generation of B6.Cg.App^tm1.1Dnli^/J (*App*^SAA^; JR#034711) knock-in mouse model is described in Xia et al. 2022 [11]. The *App*^SAA^ knock-in mouse model was engineered by insertion of 6 mutations into the genomic *App* locus via homologous recombination. Three amino acids were substituted to humanize the mouse Aβ1-42 region (APP G676R, APP F681Y and APP R684H) and three FAD-linked APP mutations were inserted: KM670/671NL (Swedish), E693G (Arctic) and T714I (Austrian). This mouse model was created and maintained on the C57BL/6J background [26]. The *App*^SAA^ mice were bred at The Jackson Laboratory (Bar Harbor, ME, United States). Age- and sex-matched C57BL/6J (JR#000664) mice were obtained from The Jackson Laboratory to serve as controls. All cohorts of mice were aged in facilities at The Jackson Laboratory, group-housed within sex and genotype (n=3-5 per cage), maintained in a 12/12-hour light/dark cycle, and room temperatures were maintained at 18-24°C (65-75°F) with 40-60% humidity. All mice were housed in positive, individually ventilated cages. Mice received ad libitum autoclaved 5K52 6% fat diet (Lab Diet, St. Louis, MO, United States) and water with acidity regulated from pH 2.5-3.0. The number of animals per sex per cohort for each experiment are listed in **Supplemental Table 1**.

### 2.2. Tissue collection

Animals were deeply anesthetized via intraperitoneal (i.p.) injection of 4% Avertin (Tribomoethanol) at 800mg/kg. Blood samples were collected via cardiac puncture in an EDTA tube (FisherScientific, Waltham, MA USA 02451; #02-669-33) then centrifuged at 12,500 rpm for 15 minutes at 4°C. The supernatant was then collected and stored at - 80°C. The mice were then transcardially perfused with cold phosphate buffered saline (PBS). The brain was carefully removed from the skull and divided midsagitally into left and right hemispheres using a brain matrix. Hemibrains were flash frozen and stored at -80°C for biochemical analyses or postfixed in 4% paraformaldehyde (PFA) overnight at 4°C for immunofluorescence followed by 15% sucrose for 24 hours at 4°C, followed by 30% sucrose for 24 hours at 4°C before storage at -80°C.

For 3D Neuropathology, mice were administered 10mg/kg of methoxy-X04 (BioTechne, Minneapolis, MN 55413; # 4920) by intraperitoneal (i.p.) injection 18-24 hours before harvest along with tail vein injections of 100μL injections of DyLight594 (1mg/mL, Vector Laboratories, Newark, CA 94560; #DL-1177-1) five minutes prior to harvest to label brain vasculature. For 3D neuropathology, mice were transcardially perfused with PBS followed by 4% paraformaldehyde diluted in PBS before removal of the brain. Whole brains were post-fixed in 4% PFA for 24 hours and then stored in 0.1% sodium azide (NaN3) until sectioning.

### 2.3. 3D Neuropathology

#### 2.3.1. Whole brain imaging

Whole brain fluorescence imaging was performed as previously described [27] with serial two-photon (STP) tomography using a TissueCyte 1000 (TissueVision Inc., Somerville, MA) with 925 nm excitation, a 500 nm dichroic mirror, and a 447/60 bandpass emission filter on the blue channel. Approximately one hundred and forty serial block-face images were acquired with a 50μm sectioning interval from each brain at 0.72 pixels/μm lateral resolution. Automated segmentation of the fluorescent signal from methoxy-X04 labeled plaques and registration to the 3D Allen Mouse Brain Common Coordinate Framework, v3 (CCFv3) [28] were performed as previously described [29]. Briefly, segmented fluorescence output is a full resolution mask that classifies each 1.38 μm × 1.38 μm pixel as either signal or background. An isotropic 3D summary of each brain is constructed by dividing each image series into 10μm × 10μm × 10μm grid voxels. Total signal is computed for each voxel by summing the number of signal positive pixels in that voxel. The 3D Allen Mouse Brain CCFv3 was aligned to each 3D image in a multistep process using both global affine and local deformable registration. Plaque density for each structure in the reference atlas ontology was calculated by summing voxels from the same structure. Plaque density in each Allen CCF brain structure was computed by ratio of total voxels representing plaque signals to total voxels of the Allen brain region. Due to the 100μm z-sampling interval, our resolution limit for detecting separate plaques in the z-axis was 100μm.

#### 2.3.2. Whole brain plaque, vessel, and co-localization quantification

Plaque quantification for Allen Mouse Atlas defined regions is reported in **Supplemental Table 2**. All image series were subjected to manual QC checks for completeness and uniformity of raw fluorescence images, minimum fluorescence intensity, and artifacts. Automatic plaque segmentation was performed using a two-fold approach. First, using an outlier-based segmentation approach to extract a mask of pixels that have an intensity greater than (Mean + Standard Deviation) the image from channel 3. Then we perform a laplacian of gaussian operation to highlight irregular rounded structures (plaques) and extract a mask of all plaque-like structures present. We then combine the two masks to extract only those plaque-like structures that have a high percentage overlap with the outlier mask. Automatic segmentation results were checked for overall quality and false positive signals by overlaying segmentation results for 3–5 single coronal sections with raw fluorescent images from STP imaging.

DyLight stained vessels are extracted using a two-fold approach as well. First, Frangi Vesselness is used to extract a mask of objects that resemble vessels. Then, color segmentation was performed to extract a secondary mask of DyLight signal by targeting the red hue color signatures. For co-localization of methoxy-X04 and DyLight signal, the two masks were then combined to extract Vessel-like objects expressing strong methoxy-X04 signal. Data was validated with a previously published CAA grading method [30]. Similar methods have been used to quantify CAA in other models [31]. Co-localization quantification by region, defined by the Allen Mouse Atlas, is included in **Supplemental Table 3.**

### 2.4. ELISA analysis of human Aβ with MSD 6E10 panel

For quantitative biochemical analysis of Aβ, soluble and insoluble proteins were extracted from brain samples based on the protocol by Casali & Landreth, 2016 [32]. Hemibrains were homogenized at 100mg/mL in tissue homogenization buffer (THB; 2mM Tris-HCl pH 7.4, 250mM sucrose, 0.5mM EDTA, 0.5mM EGTA). Soluble proteins were extracted using 0.4% DEA (diethanolamine) and insoluble proteins were extracted with formic acid. Aβ species were quantified in soluble and insoluble brain extracts in duplicate using the highly sensitive V-PLEX Aβ Peptide Panel I (K15200E) from MesoScale Discovery (MSD, Gaithsburg, MD, United States) according to manufacturer’s instructions. Data is represented as picograms of amyloid per milligram of brain tissue.

### 2.5. IBEX Multiplex immunohistochemistry

Multiplex immunohistochemistry was performed using iterative bleaching extends multiplexity (IBEX) technique [33]. Coronal 25µm sections were collected and stored floating in 31.25% glycerol (Sigma-Aldrich, G5516-1L) and 31.25% ethylene glycol (Sigma-Aldrich, 102466) dissolved in PBS at 4°C. Tissues were mounted on slides treated with chrome alum-gelatin adhesive (NewComerSupply, #1033A, Middleton, WI) before continuing with immunofluorescent labeling as previously described [30] with antibodies for each cycle of IBEX listed in **Supplemental Table 4**. After imaging using the Leica THUNDER system, fluorophores were bleached with LiBr_4_ (95% lithium borohydride; ThermoFisher Scientific, Waltham, MA, United States) diluted to 1mg/ml in diH_2_0 and further stained. Slides were bleached twice, allowing three cycles of antibodies according to **Table 1**. Images were overlayed using iMARIS software (V.11.0.1; Oxford Instruments).

### 2.6. Transcriptomics

#### 2.6.1. RNA isolation, preparation, and sequencing

Total RNA was isolated from tissue using the NucleoMag RNA Kit (Macherey-Nagel; Düren, Germany) and the KingFisher Flex purification system (ThermoFisher). Tissues were homogenized in MR1 buffer (Macherey-Nagel) using a Bead Ruptor Elite (Omni International, Bedford, NH, United States). RNA isolation was performed according to the manufacturer’s protocol. RNA concentration and quality were assessed using the Nanodrop 8000 spectrophotometer (Thermo Scientific) and the RNA ScreenTape Assay (Agilent Technologies, Santa Clara, CA, United States). Stranded libraries were constructed using the KAPA mRNA HyperPrep Kit (Roche Sequencing and Life Science, Basel, Switzerland), according to the manufacturer’s protocol. Briefly, the protocol entails isolation of polyA containing mRNA using oligo-dT magnetic beads, RNA fragmentation, first and second strand cDNA synthesis, ligation of Illumina-specific adapters containing a unique barcode sequence for each library, and PCR amplification. The quality and concentration of the libraries were assessed using the D5000 ScreenTape (Agilent Technologies) and Qubit dsDNA HS Assay (ThermoFisher), respectively, according to the manufacturers’ instructions. Libraries were pooled and sequenced by the Genome Technologies core facility at The Jackson Laboratory. All samples were sequenced 150 bp paired-end on an Illumina NovaSeq X Plus using the 10B Reagent Kit (Illumina, San Diego, CA, United States), targeting 30 million read pairs per sample. Once the data was received the samples were concatenated to have a single file for paired-end analysis.

#### 2.6.2. RNA-Sequencing data processing

RNA-Seq data were processed using nf-core/rnaseq pipeline (https://doi.org/10.5281/zenodo.1400710), using the STAR-RSEM alignment/quantification workflow (--aligner star_rsem). Reads were mapped to the mouse genome (GRCm38/mm10) using STAR [34] and gene expression was quantified with RSEM [35]. Differentially expressed genes were identified using the R Bioconductor package DESeq2 (v1.16.1) [36], with genes considered significant at a Benjamini-Hochberg adjusted p-value < 0.05.

#### 2.6.3. Human AMP-AD gene co-expression modules

Wan et al. (2020) [37] identified 30 human brain co-expression modules through a meta-analysis of differential gene expression across seven brain regions in postmortem samples from three independent late-onset Alzheimer’s disease (LOAD) cohorts. These modules were grouped into five consensus clusters [37] representing shared AD-related changes across studies and brain regions. Reactome pathway enrichment analysis was used to annotate each cluster with distinct biological themes, with pathways ranked by Bonferroni-corrected p-values.

Data for the 30 human brain co-expression modules generated by the Accelerating Medicines Partnership for Alzheimer’s Disease (AMP-AD) consortium were obtained from the Synapse data repository (https://www.synapse.org/#!Synapse:syn11932957/tables/;). The log2 fold-change values for genes within these modules were originally computed from sex-regressed, residualized counts. To enable direct comparisons between female mice and female AD cases, and between male mice and male AD cases, we replaced these sex-regressed log2 fold-change values with sex-specific values for each gene module. Case-control log2 fold-change values for all quantified genes, computed separately for males and females, were obtained from the Synapse data repository (https://www.synapse.org/Synapse:syn30821563), and used to generate distinct male-specific and female-specific datasets for the 30 modules.

#### 2.6.4. Mouse-human correlation analysis

We assessed the similarity between mouse and human disease-related expression changes by computing Pearson correlations between log2 fold change values for human AD cases versus controls and the corresponding log2 fold change values for mouse models versus controls, stratified by sex. Correlations were calculated across the set of orthologous genes within each AMP-AD module [38, 39], and across orthologous genes within corresponding gene modules using cor.test function in R as:

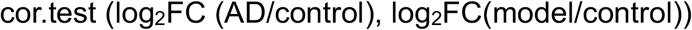

where logFC(AD/control) is the log fold change in gene expression of human AD patients compared to control patients, and log2FC (model/control) is the corresponding value in mouse models compared to controls.

Correlation results were plotted using the ggplot2 package in R. Framed circles denote significant (p < 0.05) Pearson’s correlation coefficients, with blue indicating positive and red indicating negative correlations. Both circle size and color intensity are proportional to the magnitude of the correlation coefficient.

### 2.7. Plasma Biomarkers

Mouse NULISA™ (Alamar Biosciences, Fremont, CA, United States) was used to obtain relative quantification of 120 protein analytes. Blood was drawn from mice ranging from 4- to 16-months-of-age and approximately 35μL of plasma per sample was shipped to Emtherapro, Inc. (Atlanta, GA, USA) and assayed with the NULISA™ Mouse Panel 120. Normalized protein quantification (NPQ) units were used for protein abundances following standard NULISA™ processing. All proteins passed plate-specific limits of detection in all samples. NPQ values for each analyte were fit to a linear model as:

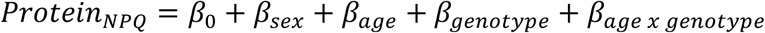

where *β*_0_ is the intercept, *β_sex_* is the effect of male sex, *β_age_* is the effect of age minus the population mean age, *β_genotype_* is the effect of the SAA genotype, and *β_a_*_g*e*_ *_x_* _g*enotype*_is the interaction term motivated by inspection of the data. Effect sizes were estimated with least squares regression and nominal p-values were reported.

NPQ values for each sample are reported in **Supplemental Table 5** and regression results for each protein are in **Supplemental Table 6**.

### 2.8. Context-dependent associative memory

Mice were handled by an experimenter for 2 minutes for at least three consecutive days prior to the extinction paradigm. Encoding and extinction was performed in Lafayette Instruments (Lafayette, IN, United States) mouse boxes (interior dimensions: 45.75 x 45.75 x 55.1 cm) equipped with cameras (3.75 frames/sec) and analyzed with FreezeFrame software (Lafayette, IN, United States). To separate the associative memory context from the extinction context, each utilized unique and distinct odors, within-animal cleaning solution, internal luminance, and floors/walls. For each session, mice were allowed 2 mins of contextual investigation prior to the onset of any tone presentations. To form an associative fear memory, mice were placed in context A and five 10 s tones (4 kHz, 70-80 dB) were paired with a 1 second, 0.8mA foot shock. For extinction sessions in context B, a series of 12 randomly dispersed 20 s 4 kHz tones were played across a 20-minute session. For extinction memory retrieval (in context B) and fear renewal (in context B), 4 tones were played during a 6-minute session. FreezeFrame immobility rates were exported to Excel (Microsoft, WA, United States) and analyzed in Prism (GraphPad, MA, United States).

### 2.9. Synaptic reconstructions

12-month-old male B6J (n=4) and *App*^SAA^ mice (n=6) were engineered to express fluorophores in specific hippocampal pyramidal neurons using brain injections of recombinant AAVs using IACUC approved methods (#20006). Under isoflurane anesthesia, mice were placed on a heating pad and secured in a Kopf stereotactic apparatus (#1902, David Kopf Intsruments, Tujunga, CA, United States) under an attached Zeiss microscope (Discovery V20, Oberkochen, Germany). Craniotomies (<1 mm in diameter) were made using a dental drill (Foredom, CT, US), and glass pipettes (20-30 µm diameter tip) containing high-titer viral suspensions (1e^12^ gc/ml) were lowered into each target region using a Sutter micromanipulator MP285 micromanipulator (Sutter, CA, United States ). To label CA1-to-PFC neurons, injections of AAVretro-Cre were made into the mPFC (+1.8 mm A/P; 0.9 M/L; −2.5 D/V at 15° tilt) and injections of AAV2/1-flex-rev-GFP were made into area CA1 (-3.4 mm A/P; 3.5 M/L; −2.5, −2.25, and −2 D/V; with all coordinates listed as distance anterior or posterior to Bregma, distance from the midline, and depth from the pial surface). At each site, ∼50 nl of viral suspension was injected in 10 nl pulses with Nanoject II (Drummond Scientific, PA, United States), the pipette remained in place for 5 minutes, and the pipette was slowly retracted following the last site. Dissolvable vicryl sutures (Ethicon, Raritan, NJ, United States) and Vetbond (3M, St. Paul, MN, United States) were used to close the scalp. Mice were administered subcutaneous Carprofen (10 mg/kg) and Buprenorphine XR (2.5 mg/kg), topical ropivacaine (0.1%) at the incision site, and received intraperitoneal injections of saline (1mL/25g) for rehydration. After surgery, mice were placed on heating pad, monitored for 10-15 minutes until they resembled experimentally-naive mice, and returned to the animal room. Mice were examined daily for 5 days following surgery to assure that incisions were healing properly, and that normal hydration and body weight was maintained.

Post-fixed brains were cut coronally on a Vibratome at 60 µm. Slices from the temporal area of CA1 containing GFP+ neurons (e.g., -3.3 to -3.5 mm posterior) were stained with the amyloid β-sheet binding fluorophore X34 (SML1954, Sigma, MA, United States) as done previously [40], mounted, coverslipped with Vectashield (Vector Laboratories, CA, United States) and imaged using a Leica SP8 confocal microscope (Wetzlar, Germany) equipped with multiple objectives, laser lines, avalanche photomultiplier tubes, PMT/HyD photodetectors, and 40x (1.3NA) or 63X (1.4NA) objectives. For synaptic reconstructions, thin-diameter dendritic branches from each dendritic compartment (e.g., distal tuft branches, apical oblique branches, and basal branches) were identified and z-stacks were imaged with a 63x 1.4 NA objective (0.05 x 0.05 x 0.1 µm voxel size) and processed through Leica Lightning Deconvolution. The dendritic cable and individual spines were analyzed in open-source NeuronStudio software [41], which allows a semiautomated analysis of individual spines along reconstructed dendritic cables. Branch spine densities were analyzed within each compartment using a one-way ANOVA with Holm-Sidak’s multiple comparisons tests using dendritic branches as individual measures.

For immunofluorescent labeling of microglia, slices were first stained with the amyloid β-sheet binding fluorophore X34 as above before immunofluorescent labeling for IBA-1 and CLEC7A **Table 1**. Slides were imaged by Leica SP8 confocal tile scans with a 40x (1.3NA) objectives and cell counts of single- and double-labeled microglia were counted in FIJI. Quantification of IBA1-positive microglia and CLEC7A-positive DAMs was performed followed by nearest neighbor analyses in which the distances between all IBA1-positive and IBA1 and CLEC7A-positive microglia were computed and the closest neighbor was included in the analysis.

### 2.10. Data Analysis

Unless stated otherwise, data were analyzed using Prism10 statistical package (GraphPad, San Diego, CA, United States). Data was analyzed with normality tests, and according to statistical assumptions, either a one-way ANOVA with Tukey’s post-hoc test or a two-way ANOVA with Holm-Sidak’s post-hoc test. p values < 0.05 were considered statistically significant.

## 3. RESULTS

### 3.1. Age-dependent plaque pathology in *App*^SAA^ mice

Whole brain, region-specific, age-dependent plaque accumulation was quantified using the TissueVision platform as previously described [6]. In homozygous *App*^SAA^ mice, images show sparse methoxy-X04-positive plaques present at 4-months, increasing with age (**Figure 1A**). Quantification revealed plaque deposition occurs in a primarily age-dependent and linear manner from 6- to 12-months-of-age, plateauing until 19-months-of-age (**Figure 1B**). While there was a small but significant difference between male and female plaque volume in the midbrain, medulla, and pons **(Supplemental Figure 1)**, there were no sex-specific effects in the cortex, hippocampus, or cerebellum or in the entire cerebrum **(Supplemental Figure 1)**. Contrasting 3D plaque neuropathology between 4- and 19-month-old *App*^SAA^ mice can be viewed in **Supplemental Videos 1 and 2.** Age-dependent quantification data of methoxy-X04 signal by region defined by the Allen Mouse Atlas can be found in **Supplemental Table 2.** This powerful technique revealed this humanized amyloid knock-in model reliably replicates age-associated amyloid pathology.

**Figure 1.**
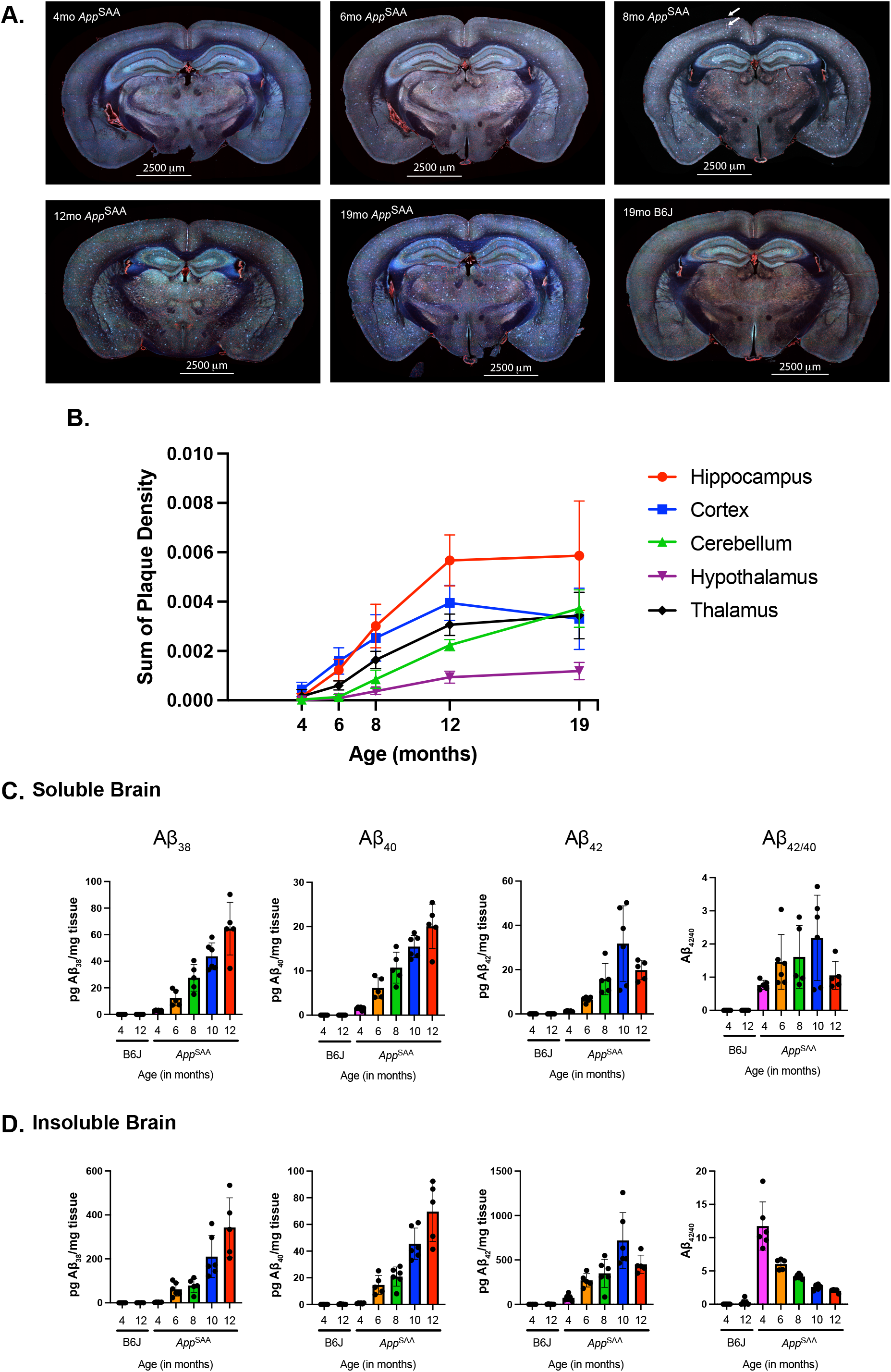
Whole brain, age-dependent plaque and amyloid accumulation in *App*^SAA^ mice. **A.** Representative whole-brain images of *App*^SAA^ mice accumulating methoxy-X04-positive plaques in an age-dependent manner from 4- to 19-months-of-age. Methoxy-X04 plaques are apparent by 4-months-of-age and largely parenchymal at younger ages. Amyloid in vasculature and leptomeninges appears by 8-months-of-age (annotated by white arrows). No methoxy-X04 signal is detected in 19-month-old B6J controls. **B.** Images were aligned to the CCFv3 Allen Brain Atlas post-methoxy-X04 labeling for whole-brain plaque quantification. Age-dependent quantitative analysis of plaques revealed a linear rate of accumulation between 4- and 12-months-of-age in *App*^SAA^ mice before levels plateau until 19-months-of-age. **C-D.** Levels of Aβ_38_, Aβ_40_, and Aβ_42_ were quantified in **C.** soluble and **D**. insoluble preparations of bulk single-hemisphere brain lysates with the MSD 6E10 panel. *App*^SAA^ mice show an age-dependent increase from 4- to 12-months-of-age in soluble and insoluble levels of Aβ_40_. Soluble and insoluble levels of Aβ_42_ followed a similar trend with a non-significant decrease from 10- to 12-months-of-age.

The plaque accumulation rates in *App*^SAA^ mice were in stark contrast to the 5xFAD model which develops significant plaque pathology by 4-months-of-age. The 5xFAD model has been previously analyzed with the same methoxy-04 imaging quantification platform from 2- to 6-months-of-age with significant sex-specific differences in plaque accumulation [6]. Both 5xFAD and *App*^SAA^ models showed significant age-dependent plaque accumulation with the highest density in the hippocampus, thalamus, and cortex (**Supplemental Figure 2**). However, the 5xFAD model showed an increase in plaque density from 2- to 6-months-of-age while *App*^SAA^ mice showed a more gradual increase from 4- to 19-months-of-age. By 6-months of age, the 5xFAD model accumulated 13-fold more plaque density in the cerebrum compared to age-matched *App*^SAA^ mice (**Supplemental Figure 2O)**. Representative brain images from the 5xFAD model can be found in Oblak et al. 2021 [6] and region-specific comparisons between the 5xFAD and *App*^SAA^ model are in **Supplemental Figure 2**.

Analysis of heterozygous *App*^SAA^ mice demonstrates a significantly delayed initiation and development of plaque pathology primarily between 12- and 19-months-of age, with nascent plaques present in the cortex by 8-months-of-age **(Supplemental Figure 3)**. These findings highlight the utility of heterozygous *App*^SAA^ knock-in mice as a model relevant to late-onset AD (LOAD), presenting novel opportunities to investigate factors contributing to disease pathologies in an age- and dose-dependent manner.

As a semi-quantitative measure of cerebral amyloid angiopathy (CAA) [31], a common co-pathology [42] of AD, we quantified whole-brain co-localization of methoxy-X04-positive plaques with DyLight594 labeled cerebral vasculature. By 8-months-of-age leptomeningeal CAA pathology is present along with strong banding of methoxy-X04 along descending arterioles (**Figure 1** indicated by arrows). While female *App*^SAA^ mice showed an age-dependent increase in co-localization of plaque and vessel signal primarily from 8- to 12-months that plateaued until 19-months, male *App*^SAA^ showed a linear increase from 6- to 19-months-of-age (**Supplemental Figure 4**). In contrast, male and female mice accumulate methoxy-X04 plaques at a similar rate (**Supplemental Figure 1**). Lastly, heterozygous *App*^SAA^ mice develop limited CAA by 19-months-of-age compared to homozygous *App*^SAA^ mice (**Supplemental Figure 5**). Data for co-localization quantification separated by Allen Mouse Atlas defined regions can be found in **Supplemental Table 3.** These findings demonstrate the translatability of this model for replicating human-relevant vascular pathologies.

To quantify amyloid species in the brain, soluble and insoluble protein fractions were assayed with a 6E10 MSD ELISA detecting human Aβ. As expected, soluble and insoluble fractions of Aβ_40_ and Aβ_42_ were significantly higher in *App*^SAA^ mice than B6J controls (**Figure 1C-D**). In *App*^SAA^ mice, there was a steady increase in both Aβ_38_ and Aβ_40_ from 4- to 12-months-of-age. Soluble and insoluble Aβ_42_ levels increased steadily from 4- to 10-months of age with a non-significant trend from 10- to 12-months-of-age (**Figure 1C-D**). While soluble Aβ_42/40_ ratio increased slightly from 4- to 10-months-of-age, 12-month ratios were similar to 4-month *App*^SAA^ mice. The insoluble Aβ_42/40_ ratio decreased steadily from a peak of 11.7 (mean) at 4 months to 2.03 at 12-months (**Figure 1C-D**). These findings further demonstrate the reliability and utility of *App*^SAA^ knock-in mice.

### 3.2. Age-dependent amyloid-associated neuropathology in *App*^SAA^ mice

Common plaque-associated proteins were evaluated using IBEX, a method of multiplexed immunofluorescence [33], on brain sections from *App*^SAA^ mice aged 4- to 12-months-of-age. Antibodies and their targets are listed in **Supplemental Table 4**. Representative images (**Figure 2**) from *App*^SAA^ mice aged 4- to 12-months-of-age revealed LAMP1-positive dystrophic neurites accumulate from 4-months of age with LAMP1 present encircling early methoxy-X04 dense core plaques with 6E10-positive diffuse halos. LAMP1 signal increased with increasing amyloid (6E10 and methoxy-X04 staining) in both cortex and hippocampal regions (**Figure 2**). Similarly, GFAP and IBA-1 increased in proximity to plaques over time. These results highlight the complex neuropathogenic processes replicated in this model, as well as identifies stages of progression of disease-associated pathology and neuroinflammation in the *App*^SAA^ model that could be targeted for therapeutic intervention.

**Figure 2.**
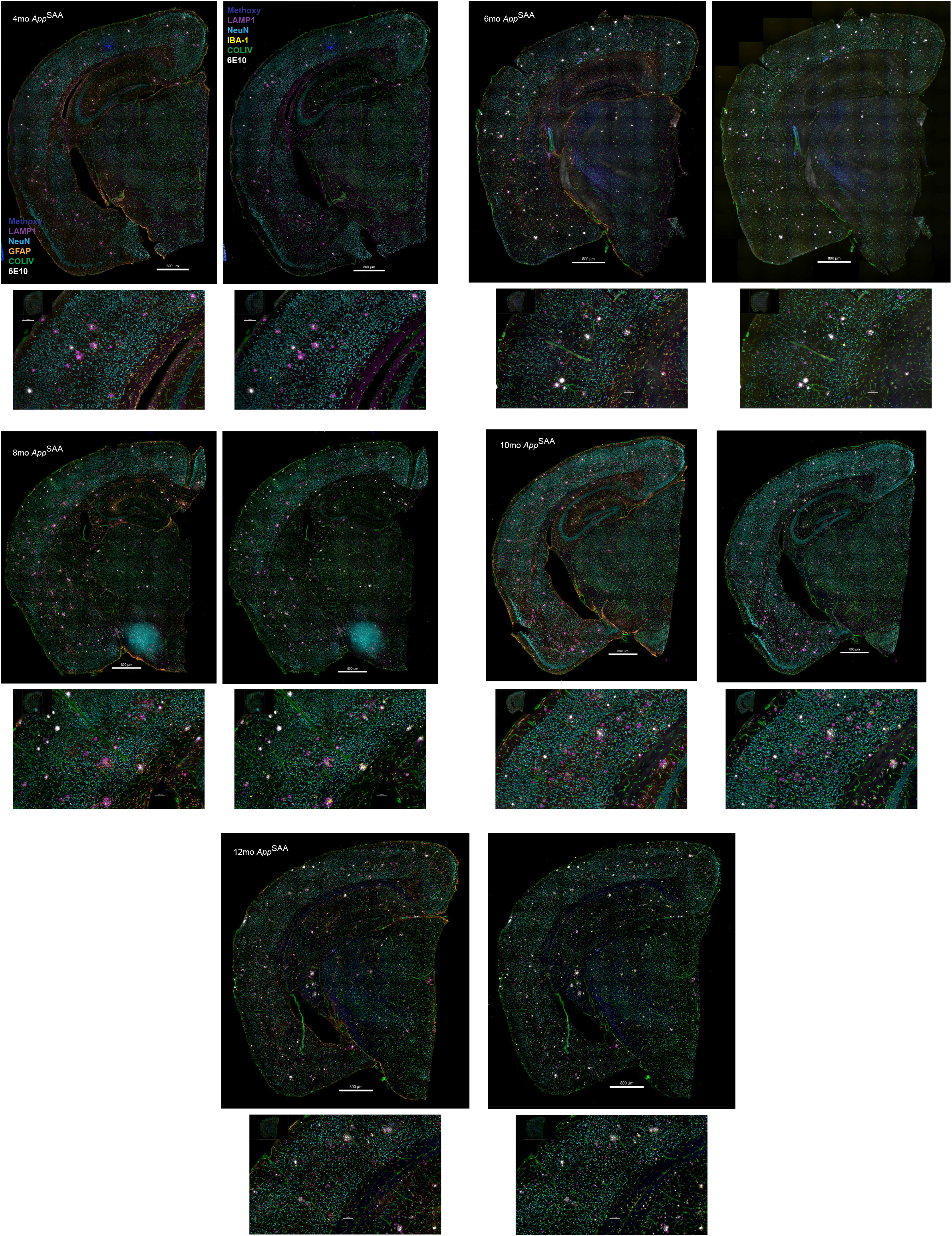
Multiplex IHC reveals age-dependent amyloid-associated pathology. Representative images of whole brain cortical brain regions imaged after multiplex immunofluorescent staining using the IBEX technique. Brains were stained *in vivo* with methoxy-X04 followed by immunofluorescent staining with antibodies for LAMP1, NeuN, GFAP, IBA-1, COLIV, and 6E10 for amyloid in 4- to 12-month *App*^SAA^ mice. LAMP1, GFAP, and IBA-1 staining is apparent surrounding methoxy-X04 and 6E10 positive amyloid plaques at 4-months of age and increases to 12-months-of-age.

### 3.3. *App*^SAA^ transcriptomic signature aligns with AMP-AD modules at 12-months-of age

We evaluated gene expression changes in hemi-brain samples from male and female *App*^SAA^ mice aged 4- and 12-months. At 4-months-of-age there were no differentially expressed genes (DEGs) detected between *App*^SAA^ mice and age-matched B6J controls. At 12-months-of-age there were over 300 DEGs (females: 342 up, 18 down; males: 398 up, 33 down). At 12-months-of age there are a number of disease-associated microglia (DAM) -related DEGs identified by bulk RNA-sequencing such as Cst7, Itgax, Ccl6, and Lyz2 (**Figure 3A**). The PCA plot highlights the separation between males and females from 4- to 12-months-of-age in both B6J and *App*^SAA^ mice. By 12-months-of-age the *App*^SAA^ mice show a transcriptomic shift away from B6J controls and 4-month *App*^SAA^ mice (**Figure 3B**). When looking at transcriptomic changes in males compared to females at 12-months, there were minimal differences **(Supplemental Figure 6A-B)**.

**Figure 3.**
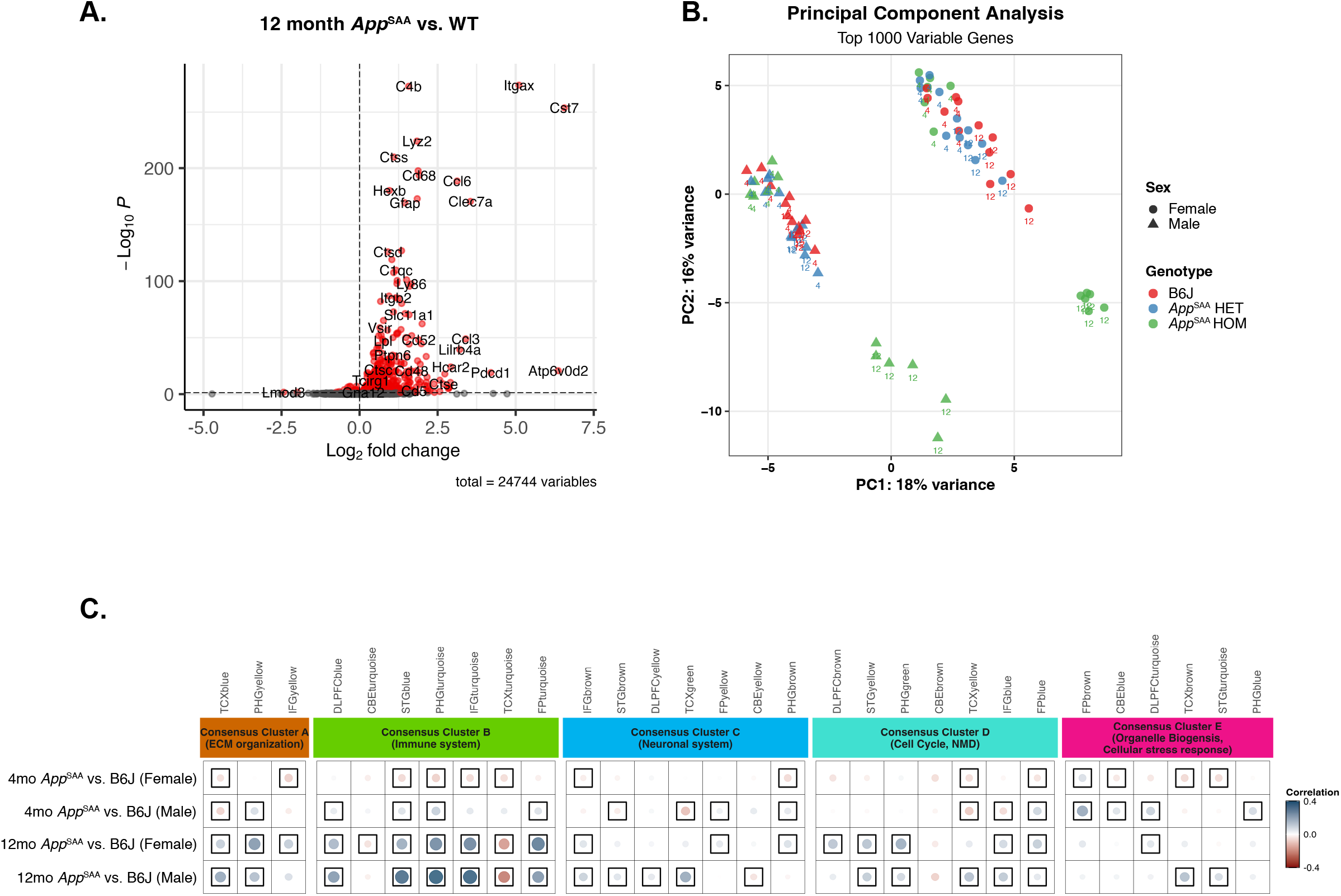
Transcriptomics from brain homogenate show an age-dependent increase in alignment to AMP-AD consensus modules from 4- to 12-months-of-age in *App*^SAA^ mice. **A.** Volcano plot highlighting significantly upregulated DEGs in 12-month *App*^SAA^ mice including previously identified microglia-specific genes. All comparisons are to age-matched B6J cohorts. **B.** The PCA plot compares sex on the x-axis to age on the y-axis, highlighting the transcriptomic shift in the 12-month homozygous *App*^SAA^ mice away from 4-month homozygous *App*^SAA^ mice and 4- and 12-month-old heterozygous *App*^SAA^ and B6J control mice. **C.** Pearson correlation coefficients for gene expression changes in mice and human disease (log fold change for cases minus controls). The five AMP-AD consensus clusters define the major functional groups of AD-related alterations. Positive correlations are shown in blue and negative correlations in red with color intensity and size of the circles proportional to the correlation coefficient. Boxes indicate statistical significance at p ≤ 0.05.

In comparison to human patient data represented in the Accelerating Medicines Partnership for AD (AMP-AD) program [39], there is little alignment with homozygous *App*^SAA^ mice at 4-months-of-age (**Figure 3C**). However, by 12-months-of age they show strong alignment with AMP-AD consensus clusters A and B (ECM organization and immune system, respectively), with weaker, but still significant associations in consensus cluster C (neuronal system) and D (cell cycle) (**Figure 3C**). Heterozygous *App*^SAA^ mice at 4- and 12-months-of age show weak alignment in non-immune AMP-AD modules **(Supplemental Figure 6C).** At 4-months-of-age the 5xFAD transgenic mouse model shows strong correlations in neuroinflammatory modules (Consensus Group B) and extracellular matrix modules (Consensus Group A) that are not significantly altered at 8 or 12 months. In summary, homozygous *App*^SAA^ mice exhibit minimal correlation at 4-months-of-age, but similar correlations as the 5xFAD model at 12-months-of-age (**Supplemental Figure 6C**). These results show the translationally-relevant development of AD-related neuropathology in the *App*^SAA^ model compared to the 5xFAD model.

### 3.4. Age-dependent plasma biomarker changes in *App*^SAA^ mice

We assayed plasma biomarkers using the Alamar NULISA™ Mouse Panel 120 to quantify key markers of neurodegenerative disorders and associated cytokines and chemokines. Cross-sectional plasma samples drawn spanning ages 4-months to 16-months were assayed to capture age-dependent changes in *App*^SAA^ and B6J control mice (**Supplemental Table 1**). Data inspection revealed numerous proteins with age-dependent changes, therefore an age-by-genotype effect was included in a linear model analysis to identify proteins modified in *App*^SAA^ mice at advanced ages.

Differential abundances were determined by linear model effect estimates (Methods; **Supplemental Table 6**). Thirty-six proteins were differentially abundant in male versus female mice (**Supplemental Figure 7A**), with the greatest effect on prolactin (PRL) (*β_sex_* = -4.2; *p* = 1.6 x 10^-18^). Among key dementia markers, TREM2 was elevated in males (*β_sex_* = 0.6; *p* = 1.0 x 10^-5^). There were only nine proteins associated with age (**Supplemental Figure 7B**) but 37 associated with age-by-genotype (**Figure 4A**), providing evidence that the majority of age-related changes in these biomarkers are driven by the presence of humanized Aβ accumulation. Although we also observed 39 proteins associated with the SAA genotype (**Supplemental Figure 7C**), genotype and age-by-genotype effects were highly correlated (ρ = 0.66, *p* = 6.7 x 10^-16^; **Supplemental Figure 7D**) and there was no evidence of proteins with age-independent genotype effects.

**Figure 4.**
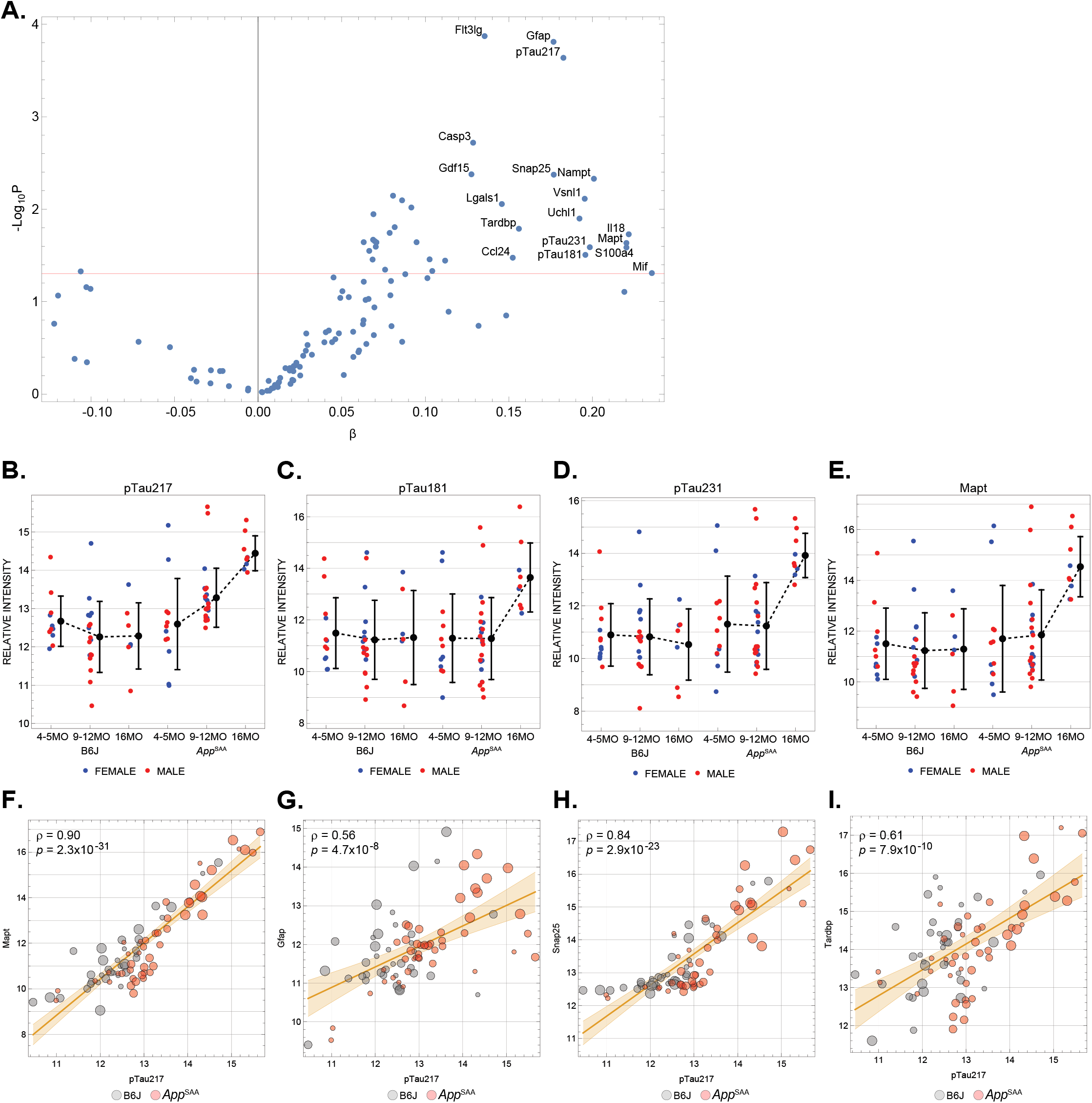
Multiple plasma biomarkers of neurodegeneration increase with age in *App*^SAA^ mice. **A**. Age-by-genotype interaction effects for Alamar NULISA biomarker panel, with positive effects corresponding to age-dependent increases in *App*^SAA^ mice. All Tau forms exhibit significant interaction. (**B-E**) Age-stratified NULISA intensities of Tau species in *App*^SAA^ mice and C57BL/6J (B6J) controls. pTau-217 is increased earliest (9-12 months, **B**) and all markers are elevated at 16 months. Black dots denote group means and error bars represent standard deviations. (**F-I**) Correlations of key biomarkers with pTau-217 levels across all study animals, showing coordinated multi-biomarker signatures across the population. Dot size denotes animal age.

Focusing on plasma protein change in aging *App*^SAA^ mice, we observed multiple pathology-related analytes increasing with age (**Figure 4A**). All four Tau species on the panel increased, with pTau-217 rising by 9-12 months (**Figure 4B**) while pTau-181, pTau-231, and total Tau (Mapt) showed increases by 16 months (**Figure 4C-E**). This Tau signature was complemented by similar age-by-SAA increases in neurodegeneration biomarkers Gfap (*β* = 0.18; *p* = 1.5 x 10^-4^), Snap25 (*β* = 0.18; *p* = 4.2 x 10^-3^), Casp3 (*β* = 0.13; *p* = 1.9 x 10^-3^), and Flt3lg (*β* = 0.14; *p* = 1.3 x 10^-4^). We also observed an increase in plasma TDP-43 (Tardbp; *β* = 0.16; *p* = 0.016). To determine if these biomarkers were similarly altered in individual animals, we assessed correlations of each biomarker across individual mice and determined that multiple age-by-genotype proteins changed similarly in each animal (**Figures 4F-I**). Correlations between Tau species were particularly strong; e.g. pTau-217 was correlated with Mapt (ρ = 0.90, *p* = 2.3 x 10^-31^; **Figure 4F**), pTau-181 (ρ = 0.79, *p* = 3.2 x 10^-19^), and pTau-231 (ρ = 0.91, *p* = 5.0 x 10^-32^). These findings suggest that amyloid accumulation drives a coordinated, robust multi-marker signature of neuropathology in *App*^SAA^ mice.

### 3.5. Memory interference in 12-month-old *App*^SAA^ mice

5xFAD and other transgenic amyloid models show memory impairments [43], so we asked whether associative memory was impaired in the *App*^SAA^ model using a context-dependent fear extinction paradigm (**Figure 5A**). We included three groups of 12-month-old mice: B6J (N=23), *App*^SAA^ (N=48), and *App*^SAA^ mice in which we omitted extinction learning (*App*^SAA:No^ ^Ext^; N=25). All mice showed strong associative learning during encoding on day one in the form of rapidly increased freezing rates after successive tone-shock pairing sessions (main effect of trial block: F (4, 438) = 113.8, P<0.001); *App*^SAA^ mice, however, showed slightly reduced freezing (main effect of group: F (1, 564) = 18.22, P<0.001; no interaction). During the first extinction learning session, there was a significant main effect of group (F (1, 276) = 24.67, P<0.001) in which the B6J mice had higher freezing rates than the *App*^SAA^ mice. We found no main effect of tone session (F (3, 276) = 2.564, P=0.0551) and no significant interaction (F (3, 276) = 0.059, P=0.98) (**Figure 5B**). B6J mice also had elevated rates during the second day of extinction learning (main effect of group: F (1, 276) = 68.65, P<0.0001), and there was a main effect of tone session (F (3, 276) = 7.595, P<0.0001) but no interaction (F (3, 276) = 0.2328, P=0.8735).

**Figure 5.**
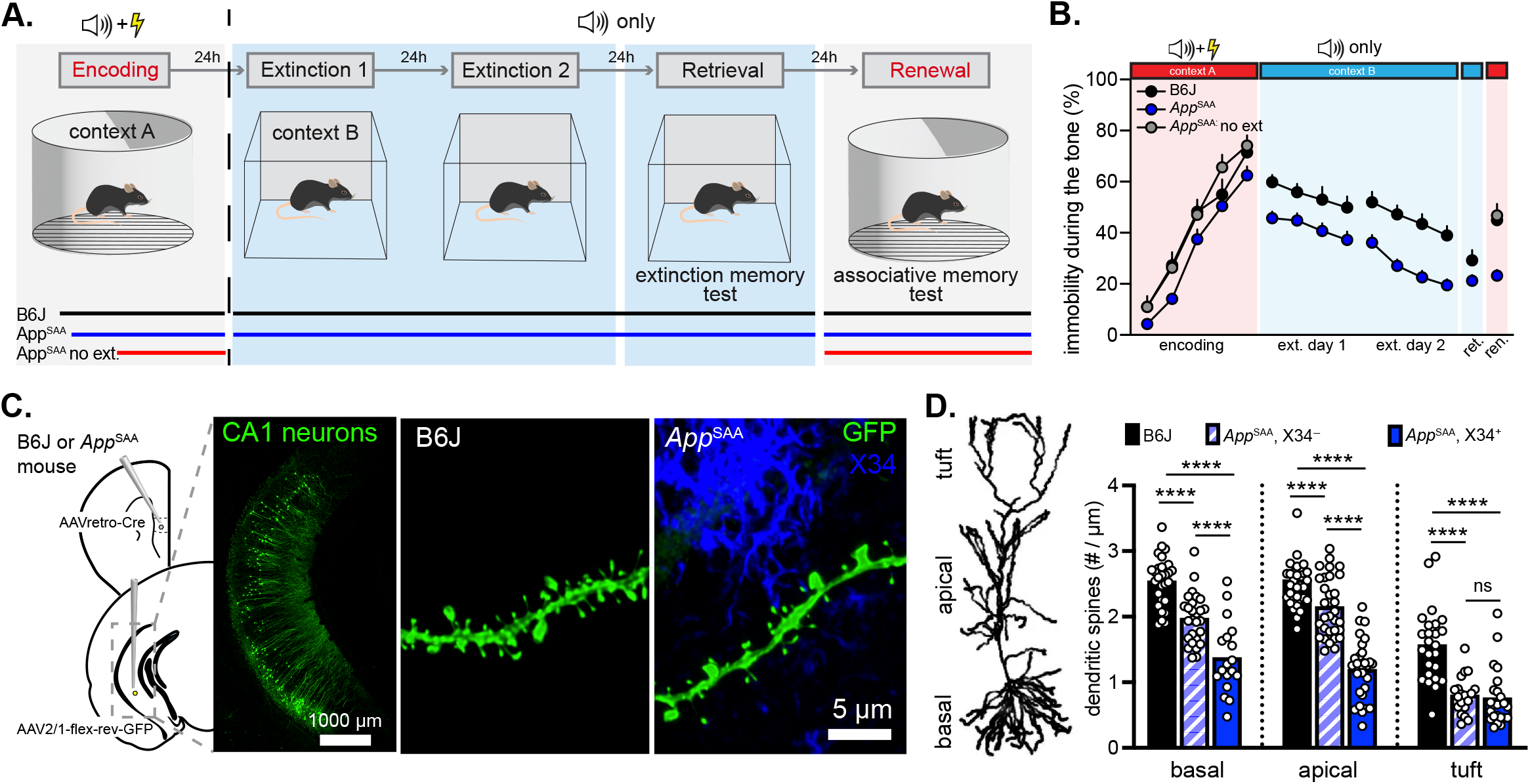
12-month-old *App*^SAA^ mice show memory deficits concomitant with spine loss. **A.** 12-month-old B6J or *App*^SAA^ mice were tested in a context-dependent extinction paradigm in which an associative auditory fear memory is learned in one context and extinguished in another. Retrieval and renewal are probe trials to determine whether mice recall the context-specific memories. **B.** B6J (black dots) but not *App*^SAA^ (blue dots) mice show recall of the associative memory in the original context. In contrast, *App*^SAA^ mice show evidence of extinction memory interference, as omitting extinction (gray dots) reveals the original memory remains intact. **C.** Labeling of CA1-to-PFC pyramidal cells in 12-month-old B6J or *App*^SAA^ mice by using AAVretro-driven GFP expression. Neurons were imaged in temporal CA1, and assigned as in proximity to plaques (i.e., within 10 µm) or in plaque-free areas. **D.** From basal, apical, and tuft dendritic compartments, branches from *App*^SAA^ mice showed reduced spine densities. Plaque proximity reduced spine density by ∼50% on apical oblique and basal branches. ****p ≤ 0.0001.

B6J and *App*^SAA^ did not differ in freezing rates during extinction memory retrieval (Mann-Whitney test; U = 418, P = 0.1). However, during the renewal task phase, where the original associative memory is recalled in a context-dependent manner, *App*^SAA^ mice froze significantly less than the B6J mice and *App*^SAA:No^ ^Ext^ mice (Kruskal-Wallis; H(3) = 27.92, P<0.0001; B6 vs. *App*^SAA^, P<0.0001; B6 vs. *App*^SAA^:no extinction, P=0.99; *App*^SAA^ vs. *App*^SAA^:no extinction, P<0.0001). Such a results shows that, in the absence of extinction learning, *App*^SAA^ mice retain the original associative memory but mice *App*^SAA^ suffer from interference between closely linked memories (**Figure 5B**).

### 3.6. Distinct localization of synapse loss on hippocampal pyramidal neurons in *App*^SAA^ mice

CA1-to-mPFC projection neurons mediate the renewal phase of the context-dependent extinction task [44] that were impaired in *App*^SAA^ mice (**Figure 5B**). To determine whether these neurons had impaired synaptic connectivity in *App*^SAA^ mice, we paired AAVretro-Cre injections into mPFC with injections of Cre-dependent AAVs encoding GFP injections into temporal CA1 to generate sparse GFP labeling of CA1-to-mPFC pyramidal neurons (**Figure 5C**). Dendritic spines were imaged on dendritic segments (n=232, 43 ± 0.7 µm in length) from 12-month-old B6J and *App*^SAA^ mice across the distal tuft, apical oblique, and basal dendritic domains (**Figure 5D**). To analyze the spatial relationship of synapses in *App*^SAA^ mice relative to plaques, dendritic branches were divided into two categories: those within 10 µm of X34 plaque material (X34+), and those located in X34 plaque-free areas (X34-).

Relative to branches from age-matched B6J mice, *App*^SAA^ mice show a reduced spine density across all three dendritic compartments. Analysis of distal tuft dendrites (B6J, n=25; *App*^SAA^:X34-, n=19, and *App*^SAA^:X34-, n=23) revealed 50% spine loss in *App*^SAA^ mice, which did not differ between X34+ and X34-branches (F(2, 64) = 23.85, P<0.0001; p<0.0001 for each *App*^SAA^ comparison to B6J branches and P=0.94 between X34+ and X34-branches). In apical oblique dendrites (B6J, n=32; *App*^SAA^:X34-, n=32, and *App*^SAA^:X34-, n=28), however, the degree of spine loss depended on plaque proximity. *App*^SAA^:X34-branches lost ∼20% of spines, while *App*^SAA^:X34-branches lost nearly 50% of spines (F(2, 89) = 60.70, P<0.0001; B6J vs. *App*^SAA^:X34-, P=0.0011, B6J vs. *App*^SAA^:X34+, P=<0.0001, and *App*^SAA^:X34- vs. *App*^SAA^:X34+, P=<0.0001). Nearly the same results were obtained from the basal compartment (B6J, n=28; *App*^SAA^ :X34-, n=27; *App*^SAA^:X34+ n=18; (F(2, 70) = 41.61, P<0.0001; B6J vs. *App*^SAA^:X34-, P=<0.0001, B6J vs. *App*^SAA^:X34+, P=<0.0001, and *App*^SAA^:X34- vs. *App*^SAA^:X34+, P=<0.0001). No X34 labeling was found in B6J mice, and dendritic spine synapse densities on dendritic branches from B6J mice were similar to previous values found using array tomography [45] or electron microscopy [46]. Such robust synapse loss at plaques was evident from neuropil staining of Synaptophysin 1, which was effectively eliminated at adjacent plaque sites **(Supplemental Figure 8A)**.

Although dendritic spine densities differ markedly between the apical oblique and distal tuft dendrites of CA1 pyramidal neurons [47], within both compartments the linear spine density scales with the parent branch diameter [47, 48]. We recapitulated this diameter-density relationship in the apical oblique and tuft branches branches from B6J mice (tuft, r= 0.55, P<0.005; apical, r=0.7, P<0.0001) and on *App*^SAA^:X34-branches (tuft, r= 0.59, P<0.01; apical, r=0.45, P<0.01; **(Supplemental Figure 8B-C**). The positive diameter-density relationship on *App*^SAA^:X34-branches presents as a similar slope with a subtractive offset, suggesting spine loss on plaque free branches occurs uniformly across branches of different diameters. However, the positive diameter-density correlation was absent in apical and tuft *App*^SAA^:X34+ branches **(Supplemental Figure 8B-C)**, suggesting that spine loss at plaques occurs in a divisive manner and is greater on higher order, thicker diameter branches **(Supplemental Figure 8B-C).** Given that branch diameter in CA1 predicts branch input impedance [49] and the expression of ion channels in the dendritic membrane [50], the subtractive (on *App*^SAA^:X34-branches) and divisive (on *App*^SAA^:X34+ branches) forms of spine loss likely have differential functional impacts on local dendritic excitability and synaptic integration.

### 3.7. DAMs are spatially positioned around plaques in the CA1 of the hippocampus

A subset of ‘disease associated microglia’ (DAM) are known to be recruited to plaques where they phagocytose amyloid and apoptotic neuronal debris [51, 52]. We evaluated whether the loss of synapses in *App*^SAA^ mice could be explained by the emergence of DAM. Relative to B6J mice, IBA1-positive microglia were more than twice as dense in the stratum pyramidale/stratum oriens (SP/SO), stratum radiatum (SR), and stratum moleculare-lacunosum (SLM) of the hippocampus of *App*^SAA^ mice **(Supplemental Figure 8D-E)**. This effect was completely accounted for by the emergence of IBA1+/CLEC7A+ putative DAM **(Supplemental Figure 8E)**. Using a nearest neighbor analysis, we found that IBA1+ microglia were spatially dispersed from each other within CA1 of B6J mice (∼40-50 µm apart from each other on average), with only ∼5% cells within 10 µm of each other **(Supplemental Figure 8F)**. This distribution was dramatically shifted in *App*^SAA^ mice, with nearly ∼40% of IBA1+ cells within 10 µm of each other. Consistent with this notion, IBA1+/CLEC7A+ microglia were essentially absent in B6J mice, and the spatial distribution of IBA1-positive, CLEC7A-negative microglia in *App*^SAA^ mice resembled that of healthy B6J mice. Thus, at plaques, the emergence and clustering of DAM render them well-positioned to mediate neuronal synapse loss in the CA1 of *App*^SAA^ mice [51, 52]. At the same time, synapse loss in the CA1 of *App*^SAA^ mice occured independent of X34 plaques (**Figure 5D**) and due to the low plaque volume in CA1 (e.g., ∼1% of the total neuropil; **Figure 1B**), our data strongly suggests that synapse loss on pyramidal neurons in CA1 primarily occurs at plaque-free dendritic locations independent of DAM. Synapse loss in the CA1 correlates with context-dependent memory impairments.

## 4. DISCUSSION

While mouse models of AD are powerful tools for hypothesis and preclinical testing, there is a need to move away from transgenic models with neuronal-specific over-expression of APP towards knock-in mouse models to improve research translatability, rigor and reproducibility [25]. With humanized Aβ under the endogenous mouse APP promoter, the *App*^SAA^ model is unaffected by artificially elevated levels of neuronal-specific human APP with mouse APP, therefore improving translatability of preclinical research. The data reported here represent a comprehensive phenotyping of neuropathology, AD-associated biomarkers, and context-dependent memory deficits defining prodromal (prophylactic) and interventional (early pathology) therapeutic windows in the *App*^SAA^ mouse model between 4- and 6-months of age and 6- and 12-months-of-age, respectively.

These therapeutic windows are reflected in 3D neuropathology showing linear, age-dependent plaque accumulation in a linear trend from 4-months until 12-months-of-age when plaque deposition plateaus until 19-months-of-age (**Figure 1A-B**). These results replicate and expand on earlier data at 2, 4, and 8 months [11], closely reflecting patterns from human post-mortem brain tissue at increasing clinical stages of amyloid accumulation [53]. During this linear phase, analysis of soluble and insoluble Aβ in bulk brain tissue also revealed an age-dependent increase in levels of Aβ_40_, with levels of Aβ_42_ following a similar pattern (**Figure 1C-D**). Multiplexed immunofluorescence confirmed age-dependent development of pathologic Aβ markers including IBA1, GFAP, and LAMP1-positive dystrophic neurites surrounding plaques (**Figure 2**), found in other transgenic and knock-in models of AD [54], as well as in post-mortem human tissue [55]. At 4-months-of-age, LAMP1 is apparent around nascent plaques in the cortex of *App*^SAA^ mice with a dense core marked by methoxy-X04 puncta and diffuse halos marked by 6E10 (**Figure 2**). This is an important hallmark of neurodegeneration found in other AD mouse models that can be used to track efficacy of disease-modifying therapies. Plaque-associated neuroinflammation marked by IBA-1 and GFAP is also a critical AD-associated process, which was reflected also reflected by in brain transcriptomic data, with 12-month old *App*^SAA^ mice showing significant increase in disease-associated microglial (DAM) genes [51] (**Figure 3A**). These transcriptomic signatures identified using bulk RNA-sequencing also overlap with the microglial specific RNA-sequencing signature from 8-month-old mice reported by Xia et al. 2022 [11].

Another amyloid-associated pathology, cerebral amyloid angiopathy (CAA), was quantified with whole brain co-localization of methoxy-X04 with DyLight stained vasculature. Our data (**Figure 1A**, **Supplemental Figure 4)** replicate and extend on original reporting of CAA at 8- and 16-months in *App*^SAA^ mice [11] with age-dependent analysis. Our findings revealed female *App*^SAA^ mice develop CAA pathology primarily between 8- and 12-months-of-age **(Supplemental Figure 4O)**, whereas male mice show a linear increase from 6- to 19-months-of-age **(Supplemental Figure 4P)** with a significant, albeit small, increase in cerebral CAA levels at 19-months compared to females **(Supplemental Figure 4B)**. This is not only a critical pathology for holistically evaluating anti-amyloid therapies because CAA pathology is present in 80% of AD patients [56], potentially more severe in males [57, 58] and associated with an increased risk of developing amyloid related imaging abnormalities (ARIA) after anti-Aβ immunotherapy [59, 60]. These data further emphasize the translatability of this model through clinically-relevant neuropathological phenotypes for anti-amyloid targeting antibodies.

The data from *App*^SAA^ mice are in stark contrast to the highly utilized 5xFAD transgenic model. Previous research has shown strong alignment between brain transcriptomic signatures of the 5xFAD model with clinical data in AMP-AD modules that is observed by 4-months and sustained to 12-months-of-age [6] while *App*^SAA^ mice showed a similar alignment from after 12-months (**Figure 3C**). In addition, whole brain volumetric plaque data in *App*^SAA^ mice contrast the early saturation in plaques in the 5xFAD model, developing approximately 13 times as much plaque volume by 6-months-of age with females developing more pathology than males [6] **(Supplemental Figure 2)**. Such early saturated levels of amyloid pathology limits the therapeutic window for testing interventional therapies. The APP^NLGF^ knock-in amyloid mouse model [61] is similar to the *App*^SAA^ model with Swedish, Arctic, and Iberian mutations showing similar age-dependent amyloid pathology. Compared to the 5xFAD model, the APP^NLGF^ model shows an approximately 12-fold difference in plaque accumulation in the hippocampus at 10-months [62], similar to the difference between 5xFAD and *App*^SAA^ mice at 6-months (**Supplemental Figure 2)**. Evaluation of heterozygous *App*^SAA^ mice highlight their utility as a model of late-onset AD (LOAD), due to the slower accumulation of plaques and CAA (**Supplemental Figures 3 and 5**). Heterozygous *App*^SAA^ mice also show slight alignment with AMP-AD modules at 12-months. Similarly, heterozygous APP^NLGF^ knock-in mice are considered a model of mild AD [63]. However, the APP^NLGF^ model has restrictions on use, whereas the *App*^SAA^ model is commercially available without restrictions.

While the *App*^SAA^ model has significant utility for AD research and preclinical testing, one important caveat lies with the incorporation of Arctic mutation in APP (E22G) which is known to induce morphological structural changes to Aβ [64]. The Arctic mutation is associated with altered binding affinity for PET probe Pittsburgh compound and dyes targeting Aβ [65], likely also affecting antibody affinity. In the original *App*^SAA^ report [11], Aβ was measured using the human and mouse-reactive 4G8 antibody specific for APP residues 18-23 [66]. However our data using the 6E10 antibody, specific to human APP residues 4-10, for ELISA revealed similar levels between 2-month mice in Xia et al. and 4-month *App*^SAA^ mice in the present study (**Figure 1C-D**). Moreover, the insoluble Aβ_42/40_ ratio shows a steady decline from peak levels at 4-months, similar to 2-month Aβ_42/40_ ratio in Xia et al. [11]. A study to qualitatively evaluate age-dependent accumulation of different amyloid structural isoforms (ie. Prefibrillar, protofibrils, and oligomers) is limited by available reagents and is out of scope for this project.

To further emphasize the clinically relevant phenotypes in this model, plasma biomarker data from *App*^SAA^ mice show a robust signature reflective of age-dependent amyloid accumulation that was absent in B6J controls. Consistent with human studies, pTau-217 showed the strongest association with combined age and *App*^SAA^ genotype, confirming it is a robust marker of brain amyloid accumulation [67–69]. While pTau-217 was elevated at both 9-12 months and 16 months of age, pTau-181, pTau-231 and total Tau were also elevated by 16 months (**Figures 4B-E**). These Tau markers were accompanied by known markers of synaptic loss including Snap25 [70] and markers of glial activation including Gfap. The coordinated, consistent changes in this AD-relevant suite of plasma proteins provides a robust panel of biomarkers of amyloid neuropathology in *App*^SAA^ mice. Such robust multi-marker signatures can serve as peripheral biomarkers in preclinical testing of amyloid-modifying therapeutics, including longitudinal studies. Furthermore, the age-dependent changes in these markers can enable the timing of treatments to assess efficacy in specific therapeutic windows.

We also show that *App*^SAA^ mice maintain long-term associative memories yet show a heightened susceptibility to interference among competing memories(**Figure 5A-B**). Past work has shown that memories can be retrieved in AD mice with context-dependent stimulation [71] and the APP/PS1 transgenic model displays a similar memory interference phenotype [72]. These results suggest that the memory storage mechanism may be less impacted by plaques compared to the consolidation or reconsolidation mechanisms that allow appropriate context-dependent memory retrieval. Such a deficit may be related to the robust synapse loss we found on CA1 pyramidal cell dendrites (**Figure 5C-D**), which mediate the recall of the original associative memory in our context-dependent task [44]. Spine loss on these CA1-to-mPFC projection neurons occurred at plaque-free sites where basal and apical branches lost ∼20% of their excitatory synapses and in proximity to plaques where the same branches lost ∼50% of spine synapses. Because plaques are relatively-sparse in CA1, our results show that the majority of synapses are lost at plaque-free locations this region, in contrast to the DAM-mediated synaptic loss observed in other brain regions. These two forms of synapse loss may be mechanistically distinct, as plaque sites are dominated by clusters DAM but plaque-free sites are not **(Supplemental Figure 8B-F)**. Interestingly, branches in the distal tuft compartment of CA1 pyramidal neurons, which receives input from the entorhinal cortex, lost ∼50% of synapses whether there was a plaque present or not (**Figure 5D**). In AD patients, synaptic degeneration strongly correlates with cognitive decline [73]. Such losses are likely to severely distort how the hippocampus relays task-relevant signals to downstream sites during memory retrieval [74, 75].

Collectively, this body of work defines therapeutic windows for prodromal and early intervention stages in homozygous *App*^SAA^ mice, with clinically relevant age-dependent increases in plaques and AD-associated phenotypes. This model will aid greatly in translation of novel biomarkers and diagnostic tools, interventional strategies and treatments into the clinic. Future work will use this model as the genetic backbone to add additional genetic risk factors such as MAPT variants and APOE alleles, as well as evaluate environmental factors like toxicant exposure and diet [25]. We hope researchers transition to the humanized amyloid *App*^SAA^ knock-in mouse model of AD to improve translatability of amyloid targeted therapies and AD-associated biomarkers and diagnostic tools.

## Acknowledgments

The authors are grateful to JAX Scientific Services, particularly Genome Technologies, Genotyping Services, and Clinical Chemistry services.

The results published here are in part based on data obtained from the AD Knowledge Portal (https://adknowledgeportal.org). Data generation was supported by the following NIH grants: P30AG10161, P30AG72975, R01AG15819, R01AG17917, R01AG036836, U01AG46152, U01AG61356, U01AG046139, P50 AG016574, R01 AG032990, U01AG046139, R01AG018023, U01AG006576, U01AG006786, R01AG025711, R01AG017216, R01AG003949, R01NS080820, U24NS072026, P30AG19610, U01AG046170, RF1AG057440, and U24AG061340, and the Cure PSP, Mayo and Michael J Fox foundations, Arizona Department of Health Services and the Arizona Biomedical Research Commission. We thank the participants of the Religious Order Study and Memory and Aging projects for the generous donation, the Sun Health Research Institute Brain and Body Donation Program, the Mayo Clinic Brain Bank, and the Mount Sinai/JJ Peters VA Medical Center NIH Brain and Tissue Repository. Data and analysis contributing investigators include Nilüfer Ertekin-Taner, Steven Younkin (Mayo Clinic, Jacksonville, FL), Todd Golde (University of Florida), Nathan Price (Institute for Systems Biology), David Bennett, Christopher Gaiteri (Rush University), Philip De Jager (Columbia University), Bin Zhang, Eric Schadt, Michelle Ehrlich, Vahram Haroutunian, Sam Gandy (Icahn School of Medicine at Mount Sinai), Koichi Iijima (National Center for Geriatrics and Gerontology, Japan), Scott Noggle (New York Stem Cell Foundation), Lara Mangravite (Sage Bionetworks).

## Sources of Funding

U54 AG054345 for MODEL-AD (LJS, DTG, NJR, MS, RP, GWC, ALO)

NIA R44 AG084491 for TissueVision, Inc (TR, RC, MS, NJR, DTG)

R01AG079877 for supporting JPT and EBB

## Consent Statement

Consent was not necessary for use of AMP-AD modules created with human data.

## REFERENCES

[1] 2025 Alzheimer’s disease facts and figures. Alzheimer’s & Dementia. 2025;21:e70235. DOI: 10.1002/alz.70235.

[2] Sasaguri H, Hashimoto S, Watamura N, Sato K, Takamura R, Nagata K, et al. Recent Advances in the Modeling of Alzheimer’s Disease. Frontiers in Neuroscience. 2022;Volume 16 - 2022. DOI: 10.3389/fnins.2022.807473.

[3] Oddo S, Caccamo A, Kitazawa M, Tseng BP, LaFerla FM. Amyloid deposition precedes tangle formation in a triple transgenic model of Alzheimer’s disease. Neurobiology of Aging. 2003;24:1063–70. DOI: 10.1016/j.neurobiolaging.2003.08.012.

[4] Jankowsky JL, Slunt HH, Ratovitski T, Jenkins NA, Copeland NG, Borchelt DR. Co-expression of multiple transgenes in mouse CNS: a comparison of strategies. Biomolecular Engineering. 2001;17:157–65. DOI: 10.1016/S1389-0344(01)00067-3.

[5] Oakley H, Cole SL, Logan S, Maus E, Shao P, Craft J, et al. Intraneuronal β-Amyloid Aggregates, Neurodegeneration, and Neuron Loss in Transgenic Mice with Five Familial Alzheimer’s Disease Mutations: Potential Factors in Amyloid Plaque Formation. The Journal of Neuroscience. 2006;26:10129–40. DOI: 10.1523/jneurosci.1202-06.2006.

[6] Oblak AL, Lin PB, Kotredes KP, Pandey RS, Garceau D, Williams HM, et al. Comprehensive Evaluation of the 5XFAD Mouse Model for Preclinical Testing Applications: A MODEL-AD Study. Frontiers in Aging Neuroscience. 2021;Volume 13 - 2021. DOI: 10.3389/fnagi.2021.713726.

[7] Forner S, Kawauchi S, Balderrama-Gutierrez G, Kramár EA, Matheos DP, Phan J, et al. Systematic phenotyping and characterization of the 5xFAD mouse model of Alzheimer’s disease. Sci Data. 2021;8:270. DOI: 10.1038/s41597-021-01054-y.

[8] Jiao S-S, Bu X-L, Liu Y-H, Zhu C, Wang Q-H, Shen L-L, et al. Sex Dimorphism Profile of Alzheimer’s Disease-Type Pathologies in an APP/PS1 Mouse Model. Neurotoxicity Research. 2016;29:256–66. DOI: 10.1007/s12640-015-9589-x.

[9] Watamura N, Sato K, Saido TC. Mouse models of Alzheimer’s disease for preclinical research. Neurochemistry International. 2022;158:105361. DOI: 10.1016/j.neuint.2022.105361.

[10] Banik A, Brown RE, Bamburg J, Lahiri DK, Khurana D, Friedland RP, et al. Translation of pre-clinical studies into successful clinical trials for Alzheimer’s disease: what are the roadblocks and how can they be overcome? Journal of Alzheimer’s Disease. 2015;47:815–43. DOI:

[11] Xia D, Lianoglou S, Sandmann T, Calvert M, Suh JH, Thomsen E, et al. Novel App knock-in mouse model shows key features of amyloid pathology and reveals profound metabolic dysregulation of microglia. Molecular Neurodegeneration. 2022;17:41. DOI: 10.1186/s13024-022-00547-7.

[12] de Weerd L, Hummel S, Müller SA, Paris I, Sandmann T, Eichholtz M, et al. Early intervention anti-Aβ immunotherapy attenuates microglial activation without inducing exhaustion at residual plaques. Molecular Neurodegeneration. 2025;20:92. DOI: 10.1186/s13024-025-00878-1.

[13] Foley KE, Weekman EM, Wilcock DM. Acute anti-Aβ antibody exposure induces microglial changes and significantly alters chemokine signaling. Alzheimer’s & Dementia: Translational Research & Clinical Interventions. 2026;12:e70201. DOI:

[14] van Lengerich B, Zhan L, Xia D, Chan D, Joy D, Park JI, et al. A TREM2-activating antibody with a blood-brain barrier transport vehicle enhances microglial metabolism in Alzheimer’s disease models. Nat Neurosci. 2023;26:416–29. DOI: 10.1038/s41593-022-01240-0.

[15] Princen K, Van Dooren T, van Gorsel M, Louros N, Yang X, Dumbacher M, et al. Pharmacological modulation of septins restores calcium homeostasis and is neuroprotective in models of Alzheimer’s disease. Science. 2024;384:eadd6260. DOI:

[16] Chang R, Chandrashekar DV, Roules GC, Jagadeesan N, Iwasaki E, Oyegbesan A, et al. Longitudinal pharmacokinetic and safety studies of an antibody-erythropoietin fusion protein for Alzheimer’s disease. International Journal of Pharmaceutics: X. 2025;10:100459. DOI: 10.1016/j.ijpx.2025.100459.

[17] Sosa Perez S, Urrutia Amable N, Viada González CE, Lorenzo-Luaces P, Sanchez Valdes L, Crombet Ramos T, et al. NeuroEPO plus (NeuralCIM®) in Alzheimer’s disease: Post-trial observational follow-up study. Journal of Alzheimer’s Disease. 2025;108:1422–35. DOI:

[18] Sosa S, Bringas G, Urrutia N, Peñalver AI, López D, González E, et al. NeuroEPO plus (NeuralCIM®) in mild-to-moderate Alzheimer’s clinical syndrome: the ATHENEA randomized clinical trial. Alzheimer’s Research & Therapy. 2023;15:215. DOI: 10.1186/s13195-023-01356-w.

[19] Shojaei M, Schaefer R, Schlepckow K, Kunze LH, Struebing FL, Brunner B, et al. PET imaging of microglia in Alzheimer’s disease using copper-64 labeled TREM2 antibodies. Theranostics. 2024;14:6319. DOI:

[20] Salarian M, Liu S, Tsai H-m, Leslie SN, Hayes T, Lo S-t, et al. Evaluation of [18F]JNJ-CSF1R-1 as a Positron Emission Tomography Ligand Targeting Colony-Stimulating Factor 1 Receptor. Molecular Imaging and Biology. 2025;27:163–72. DOI: 10.1007/s11307-025-01991-9.

[21] Chen J, Hadi F, Wen X, Zhao W, Xu M, Xue S, et al. Transcriptional regulation by PHGDH drives amyloid pathology in Alzheimer’s disease. Cell. 2025;188:3513–29. e26. DOI:

[22] Brustovetsky T, Khanna R, Brustovetsky N. Collapsin Response Mediator Protein 2 (CRMP2) Modulates Mitochondrial Oxidative Metabolism in Knock-In AD Mouse Model. Cells. 2025;14:647. DOI:

[23] Blackmer-Raynolds L, Lipson LD, Fraccaroli I, Krout IN, Chang J, Sampson TR. Longitudinal characterization reveals behavioral impairments in aged APP knock in mouse models. Scientific Reports. 2025;15:4631. DOI: 10.1038/s41598-025-89051-8.

[24] Macheda T, Hawkins MR, Johnson CE, Lapid MG, Whitlock HR, Shepard SM, et al. Glial cytokine modulation improves sleep and circadian disruption in female SAA knock-in mice of Alzheimer’s-related pathology. Alzheimers Dement. 2026;22:e71314. DOI: 10.1002/alz.71314.

[25] Oblak AL, Sasner M, Carter GW, Howell GR, Sukoff Rizzo SJ, Leal K, et al. The past, the present, and the future of preclinical mouse models for Alzheimer’s disease and related dementias. Alzheimer’s & Dementia. 2026;22. DOI: 10.1002/alz.71741.

[26] Koentgen F, Lin J, Katidou M, Chang I, Khan M, Watts J, et al. Exclusive transmission of the embryonic stem cell-derived genome through the mouse germline. genesis. 2016;54:326–33. DOI: 10.1002/dvg.22938.

[27] Ragan T, Kadiri LR, Venkataraju KU, Bahlmann K, Sutin J, Taranda J, et al. Serial two-photon tomography for automated ex vivo mouse brain imaging. Nature methods. 2012;9:255–8. DOI:

[28] Wang Q, Ding SL, Li Y, Royall J, Feng D, Lesnar P, et al. The Allen Mouse Brain Common Coordinate Framework: A 3D Reference Atlas. Cell. 2020;181:936–53 e20. DOI: 10.1016/j.cell.2020.04.007.

[29] Whitesell JD, Buckley AR, Knox JE, Kuan L, Graddis N, Pelos A, et al. Whole brain imaging reveals distinct spatial patterns of amyloid beta deposition in three mouse models of Alzheimer’s disease. Journal of Comparative Neurology. 2019;527:2122–45. DOI: 10.1002/cne.24555.

[30] Marola OJ, Uyar A, Keezer KJ, Chong Chie JAK, Elk KJ, Cullen AE, et al. WSB.APP/PS1 mice develop age-dependent cerebral amyloid angiopathy, cerebrovascular dysfunction, and white matter deficits. Alzheimer’s & Dementia. 2026;22:e71476. DOI: 10.1002/alz.71476.

[31] Bonnar O, Saadi F, Sanchez-Mico MV, Hanlin LH, Vom Eigen KA, Mumbi N, et al. Longitudinal multiphoton imaging of cerebral amyloid angiopathy in response to anti-ApoE4 immunotherapy in mice. Molecular Neurodegeneration. 2026. DOI: 10.1186/s13024-026-00957-x.

[32] Casali BT, Landreth GE. Aβ Extraction from Murine Brain Homogenates. Bio-protocol. 2016;6:e1787. DOI: 10.21769/BioProtoc.1787.

[33] Radtke AJ, Chu CJ, Yaniv Z, Yao L, Marr J, Beuschel RT, et al. IBEX: an iterative immunolabeling and chemical bleaching method for high-content imaging of diverse tissues. Nature Protocols. 2022;17:378–401. DOI: 10.1038/s41596-021-00644-9.

[34] Dobin A, Davis CA, Schlesinger F, Drenkow J, Zaleski C, Jha S, et al. STAR: ultrafast universal RNA-seq aligner. Bioinformatics. 2013;29:15–21. DOI: 10.1093/bioinformatics/bts635.

[35] Li B, Dewey CN. RSEM: accurate transcript quantification from RNA-Seq data with or without a reference genome. BMC Bioinformatics. 2011;12:323. DOI: 10.1186/1471-2105-12-323.

[36] Love MI, Huber W, Anders S. Moderated estimation of fold change and dispersion for RNA-seq data with DESeq2. Genome Biology. 2014;15:550. DOI: 10.1186/s13059-014-0550-8.

[37] Wan YW, Al-Ouran R, Mangleburg CG, Perumal TM, Lee TV, Allison K, et al. Meta-Analysis of the Alzheimer’s Disease Human Brain Transcriptome and Functional Dissection in Mouse Models. Cell Rep. 2020;32:107908. DOI: 10.1016/j.celrep.2020.107908.

[38] Pandey RS, Graham L, Uyar A, Preuss C, Howell GR, Carter GW. Genetic perturbations of disease risk genes in mice capture transcriptomic signatures of late-onset Alzheimer’s disease. Mol Neurodegener. 2019;14:50. DOI: 10.1186/s13024-019-0351-3.

[39] Preuss C, Pandey R, Piazza E, Fine A, Uyar A, Perumal T, et al. A novel systems biology approach to evaluate mouse models of late-onset Alzheimer’s disease. Mol Neurodegener. 2020;15:67. DOI: 10.1186/s13024-020-00412-5.

[40] Heuer SE, Keezer KJ, Hewes AA, Onos KD, Graham KC, Howell GR, et al. Control of hippocampal synaptic plasticity by microglia–dendrite interactions depends on genetic context in mouse models of Alzheimer’s disease. Alzheimer’s & Dementia. 2024;20:601–14. DOI: 10.1002/alz.13440.

[41] Rodriguez A, Ehlenberger DB, Hof PR, Wearne SL. Rayburst sampling, an algorithm for automated three-dimensional shape analysis from laser scanning microscopy images. Nat Protoc. 2006;1:2152–61. DOI: 10.1038/nprot.2006.313.

[42] Inoue Y, Shue F, Bu G, Kanekiyo T. Pathophysiology and probable etiology of cerebral small vessel disease in vascular dementia and Alzheimer’s disease. Molecular neurodegeneration. 2023;18:46. DOI:

[43] Sasaguri H, Nilsson P, Hashimoto S, Nagata K, Saito T, De Strooper B, et al. APP mouse models for Alzheimer’s disease preclinical studies. The EMBO journal. 2017;36:2473–87. DOI:

[44] Marek R, Jin J, Goode TD, Giustino TF, Wang Q, Acca GM, et al. Hippocampus-driven feed-forward inhibition of the prefrontal cortex mediates relapse of extinguished fear. Nat Neurosci. 2018;21:384–92. DOI: 10.1038/s41593-018-0073-9.

[45] Bloss EB, Cembrowski MS, Karsh B, Colonell J, Fetter RD, Spruston N. Structured Dendritic Inhibition Supports Branch-Selective Integration in CA1 Pyramidal Cells. Neuron. 2016;89:1016–30. DOI: 10.1016/j.neuron.2016.01.029.

[46] Bloss EB, Cembrowski MS, Karsh B, Colonell J, Fetter RD, Spruston N. Single excitatory axons form clustered synapses onto CA1 pyramidal cell dendrites. Nat Neurosci. 2018;21:353–63. DOI: 10.1038/s41593-018-0084-6.

[47] Megias M, Emri Z, Freund TF, Gulyas AI. Total number and distribution of inhibitory and excitatory synapses on hippocampal CA1 pyramidal cells. Neuroscience. 2001;102:527–40. DOI: 10.1016/s0306-4522(00)00496-6.

[48] Dumitriu D, Rodriguez A, Morrison JH. High-throughput, detailed, cell-specific neuroanatomy of dendritic spines using microinjection and confocal microscopy. Nat Protoc. 2011;6:1391–411. DOI: 10.1038/nprot.2011.389.

[49] Harnett MT, Makara JK, Spruston N, Kath WL, Magee JC. Synaptic amplification by dendritic spines enhances input cooperativity. Nature. 2012;491:599–602. DOI: 10.1038/nature11554.

[50] Nusser Z. Variability in the subcellular distribution of ion channels increases neuronal diversity. Trends Neurosci. 2009;32:267–74. DOI: 10.1016/j.tins.2009.01.003.

[51] Keren-Shaul H, Spinrad A, Weiner A, Matcovitch-Natan O, Dvir-Szternfeld R, Ulland TK, et al. A Unique Microglia Type Associated with Restricting Development of Alzheimer’s Disease. Cell. 2017;169:1276–90 e17. DOI: 10.1016/j.cell.2017.05.018.

[52] Hong S, Beja-Glasser VF, Nfonoyim BM, Frouin A, Li S, Ramakrishnan S, et al. Complement and microglia mediate early synapse loss in Alzheimer mouse models. Science. 2016;352:712–6. DOI: 10.1126/science.aad8373.

[53] Thal DR, Rüb U, Orantes M, Braak H. Phases of Aβ-deposition in the human brain and its relevance for the development of AD. Neurology. 2002;58:1791–800. DOI: doi:10.1212/WNL.58.12.1791.

[54] Sharoar MG, Palko S, Ge Y, Saido TC, Yan R. Accumulation of saposin in dystrophic neurites is linked to impaired lysosomal functions in Alzheimer’s disease brains. Molecular neurodegeneration. 2021;16:45. DOI:

[55] Barrachina M, Maes T, Buesa C, Ferrer I. Lysosome-associated membrane protein 1 (LAMP-1) in Alzheimer’s disease. Neuropathology and applied neurobiology. 2006;32:505–16. DOI:

[56] Biffi A, Greenberg SM. Cerebral amyloid angiopathy: a systematic review. Journal of Clinical Neurology. 2011;7:1–9. DOI:

[57] Shinohara M, Murray ME, Frank RD, Shinohara M, DeTure M, Yamazaki Y, et al. Impact of sex and APOE4 on cerebral amyloid angiopathy in Alzheimer’s disease. Acta Neuropathologica. 2016;132:225–34. DOI: 10.1007/s00401-016-1580-y.

[58] Tanskanen M, Mäkelä M, Myllykangas L, Notkola IL, Polvikoski T, Sulkava R, et al. Prevalence and severity of cerebral amyloid angiopathy: a population-based study on very elderly Finns (Vantaa 85+). Neuropathology and Applied Neurobiology. 2012;38:329–36. DOI: 10.1111/j.1365-2990.2011.01219.x.

[59] Greenberg SM, Bacskai BJ, Hernandez-Guillamon M, Pruzin J, Sperling R, van Veluw SJ. Cerebral amyloid angiopathy and Alzheimer disease — one peptide, two pathways. Nature Reviews Neurology. 2020;16:30–42. DOI: 10.1038/s41582-019-0281-2.

[60] Hampel H, Elhage A, Cho M, Apostolova LG, Nicoll JAR, Atri A. Amyloid-related imaging abnormalities (ARIA): radiological, biological and clinical characteristics. Brain. 2023;146:4414–24. DOI: 10.1093/brain/awad188.

[61] Saito T, Matsuba Y, Mihira N, Takano J, Nilsson P, Itohara S, et al. Single App knock-in mouse models of Alzheimer’s disease. Nat Neurosci. 2014;17:661–3. DOI: 10.1038/nn.3697.

[62] Locci A, Orellana H, Rodriguez G, Gottliebson M, McClarty B, Dominguez S, et al. Comparison of memory, affective behavior, and neuropathology in APPNLGF knock-in mice to 5xFAD and APP/PS1 mice. Behavioural Brain Research. 2021;404:113192. DOI: 10.1016/j.bbr.2021.113192.

[63] Milicevic KD, Abreu AC, Chandran G, Zhu YMD, Hu X, Ivanova VO, et al. Prodromal changes in cortical neuron physiology before amyloid pathology in a mild model of Alzheimer’s disease. Scientific Reports. 2026;16:16761. DOI: 10.1038/s41598-026-47370-4.

[64] Yang Y, Zhang W, Murzin AG, Schweighauser M, Huang M, Lövestam S, et al. Cryo-EM structures of amyloid-β filaments with the Arctic mutation (E22G) from human and mouse brains. Acta Neuropathologica. 2023;145:325–33. DOI: 10.1007/s00401-022-02533-1.

[65] Condello C, Lemmin T, Stöhr J, Nick M, Wu Y, Maxwell AM, et al. Structural heterogeneity and intersubject variability of Aβ in familial and sporadic Alzheimer’s disease. Proceedings of the National Academy of Sciences. 2018;115:E782–E91. DOI: doi:10.1073/pnas.1714966115.

[66] Baghallab I, Reyes-Ruiz JM, Abulnaja K, Huwait E, Glabe C. Epitomic Characterization of the Specificity of the Anti-Amyloid Aβ Monoclonal Antibodies 6E10 and 4G8. J Alzheimers Dis. 2018;66:1235–44. DOI: 10.3233/jad-180582.

[67] Benina N, Buitrago L, De Simone FI, Radwan RR, Miller MC, Martin K, et al. Plasma pTau 217/β-amyloid 1-42 ratio for enhanced accuracy and reduced uncertainty in detecting amyloid pathology. Brain. 2026;149:1168–81. DOI: 10.1093/brain/awag001.

[68] Malek-Ahmadi M, Sharma S, Stipho F, Denkinger M, Singh A, Brum WS, et al. Plasma Phosphorylated Tau 217 and Amyloid Burden in Older Adults Without Cognitive Impairment: A Meta-Analysis. JAMA Neurol. 2026;83:13–9. DOI: 10.1001/jamaneurol.2025.4721.

[69] Yu L, Boyle PA, Janelidze S, Petyuk VA, Wang T, Bennett DA, et al. Plasma p-tau181 and p-tau217 in discriminating PART, AD and other key neuropathologies in older adults. Acta Neuropathol. 2023;146:1–11. DOI: 10.1007/s00401-023-02570-4.

[70] Gaur A, Wong M, Chen JJ, Kang Y, Tahoulas D, Jeor K, et al. Synaptic biomarkers in Alzheimer’s disease dementia and mild cognitive impairment: A systematic review and meta-analysis. Alzheimers Dement. 2026;22:e71501. DOI: 10.1002/alz.71501.

[71] Roy DS, Arons A, Mitchell TI, Pignatelli M, Ryan TJ, Tonegawa S. Memory retrieval by activating engram cells in mouse models of early Alzheimer’s disease. Nature. 2016;531:508–12. DOI: 10.1038/nature17172.

[72] Poll S, Mittag M, Musacchio F, Justus LC, Giovannetti EA, Steffen J, et al. Memory trace interference impairs recall in a mouse model of Alzheimer’s disease. Nat Neurosci. 2020;23:952–8. DOI: 10.1038/s41593-020-0652-4.

[73] Tzioras M, McGeachan RI, Durrant CS, Spires-Jones TL. Synaptic degeneration in Alzheimer disease. Nature Reviews Neurology. 2023;19:19–38. DOI: 10.1038/s41582-022-00749-z.

[74] Grienberger C, Magee JC. Entorhinal cortex directs learning-related changes in CA1 representations. Nature. 2022;611:554–62. DOI: 10.1038/s41586-022-05378-6.

[75] Bittner KC, Grienberger C, Vaidya SP, Milstein AD, Macklin JJ, Suh J, et al. Conjunctive input processing drives feature selectivity in hippocampal CA1 neurons. Nat Neurosci. 2015;18:1133–42. DOI: 10.1038/nn.4062.

