## Supplemental Figures for "Characterization of the knock-in *App*^SAA^ mouse model as a powerful tool for Alzheimer’s disease preclinical research"

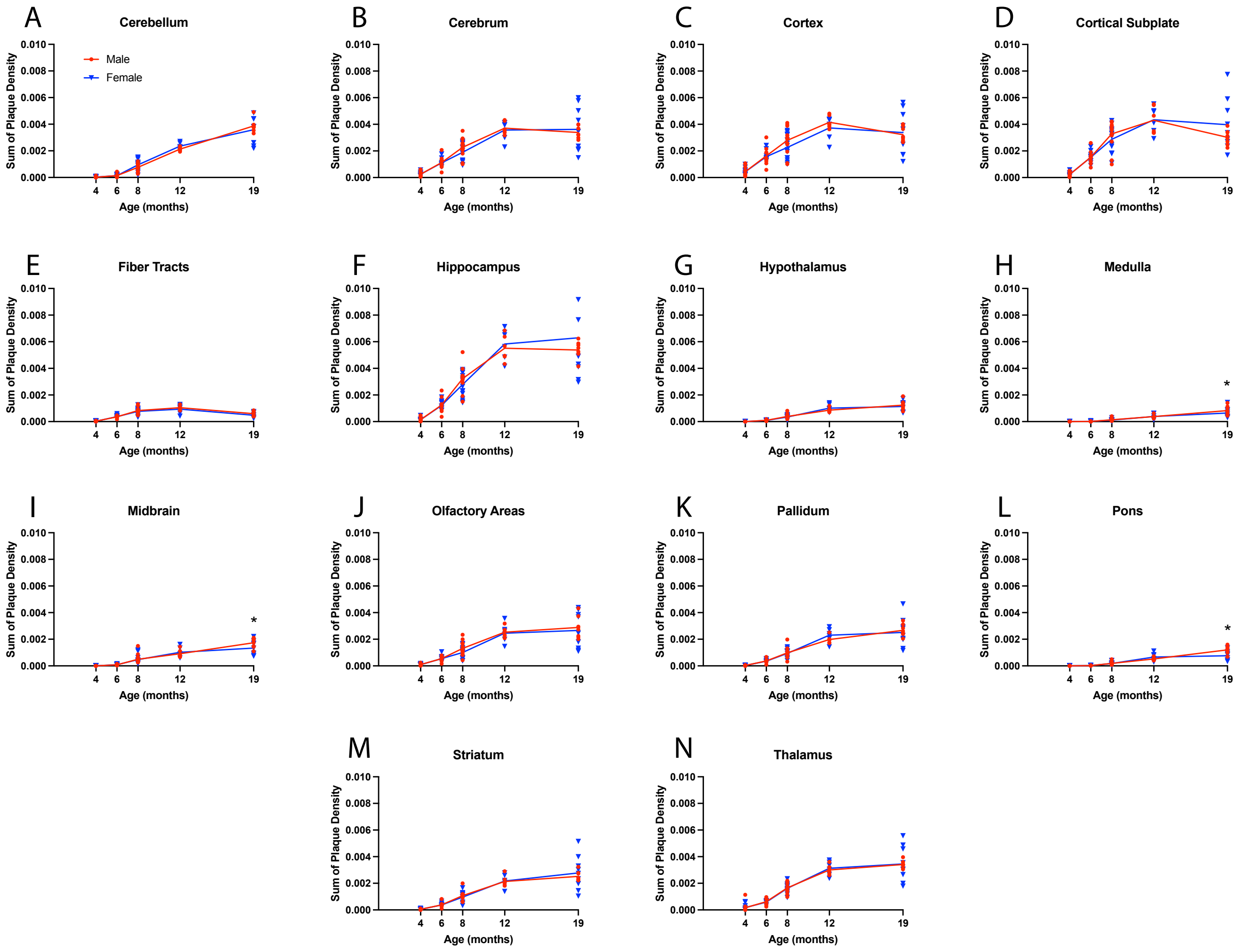


**Supplemental Figure 1. A-N.** Sex-specific analyses of methoxy-X04 plaque density in *App*$^{SAA}$ mice in the 13 major brain regions and the cerebrum, according to the Allen Brain Atlas. Sex-specific differences were small, but significant only at 19-months-of-age in the medulla, midbrain, and pons with males showing higher density of plaques compared to females. * $p \leq 0.05$ using a two-way ANOVA with Tukey's test for multiple comparisons.

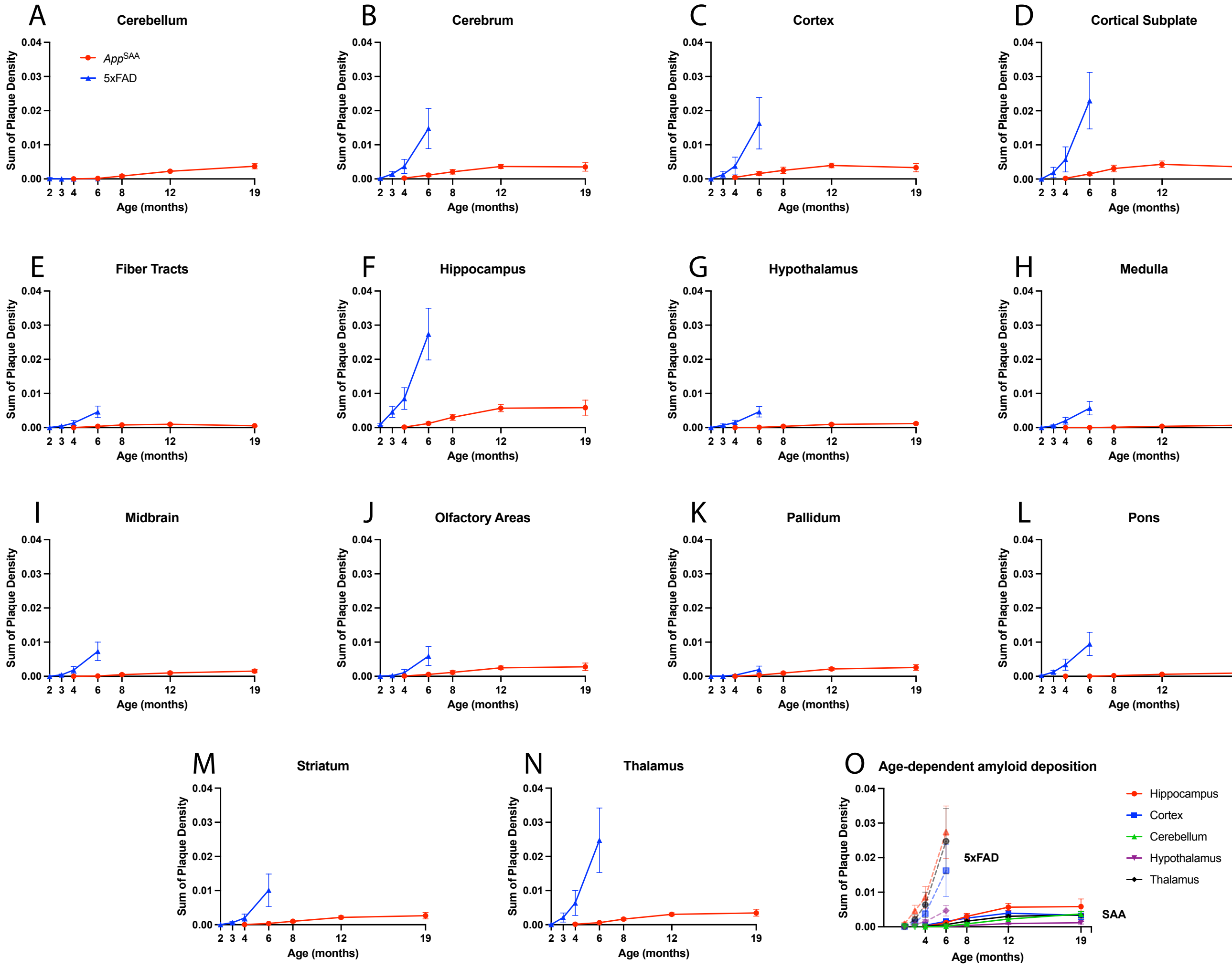


**Supplemental Figure 2. A-N.** Model comparison of methoxy-X04 plaque density between *App*$^{SAA}$ knock-in mice and 5xFAD transgenic mice aged 2- to 6-months-of-age in the 13 major brain regions and the cerebrum, according to the Allen Brain Atlas. Both models accumulate the highest plaque density in the hippocampus, thalamus, and cortex. **O.** In comparison to the linear increase from 4- to 12-months of age in *App*$^{SAA}$ mice, 5xFAD mice show rapid accumulation of plaque pathology from 2- to 6-months of age.

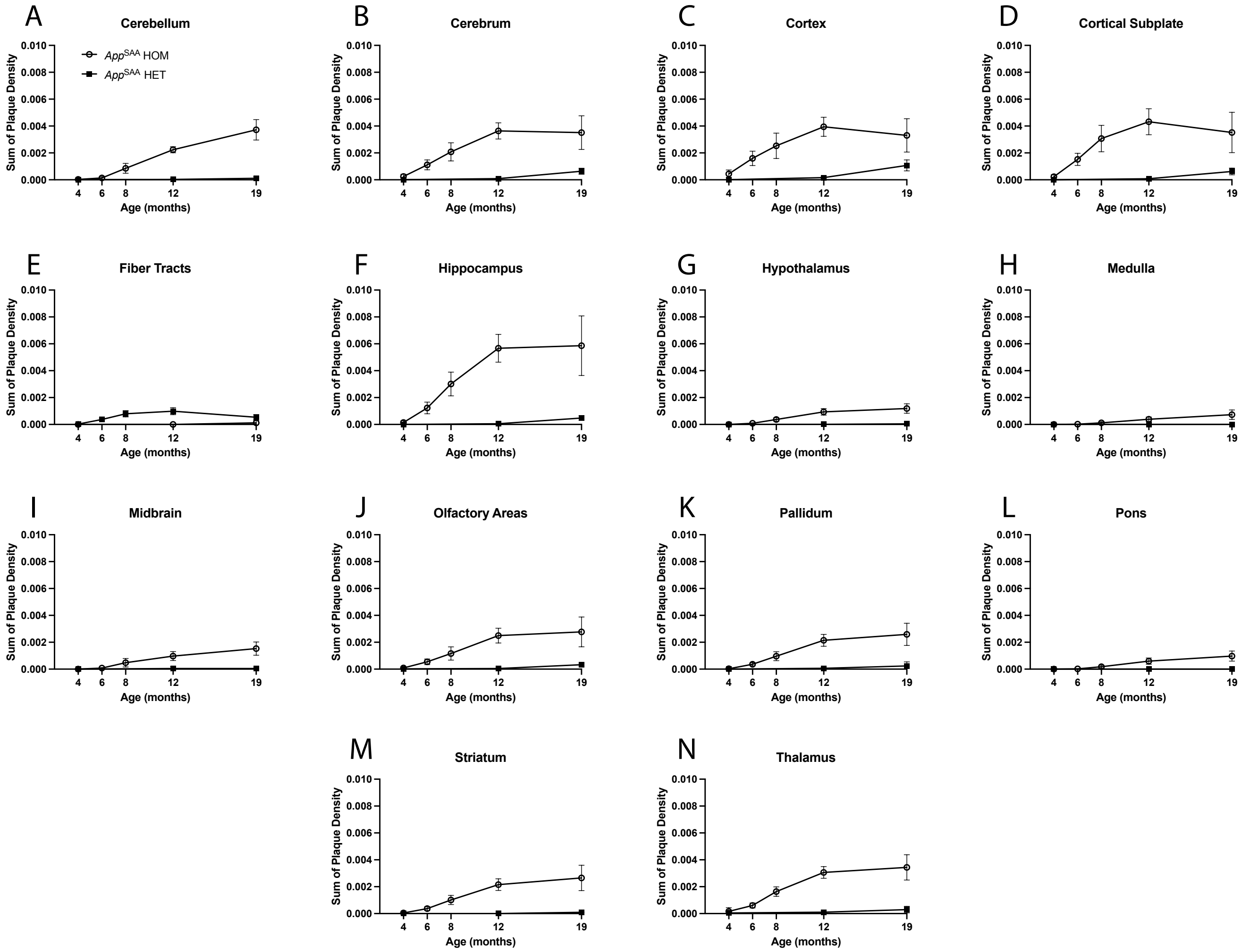


**Supplemental Figure 3. A-N.** Genotype comparisons between homozygous and heterozygous *App*$^{SAA}$ mice in the 13 major brain regions and the cerebrum, according to the Allen Brain Atlas. Heterozygous *App*$^{SAA}$ mice show minor plaque density accumulation in a region-specific manner between 12- and 19-months-of-age.

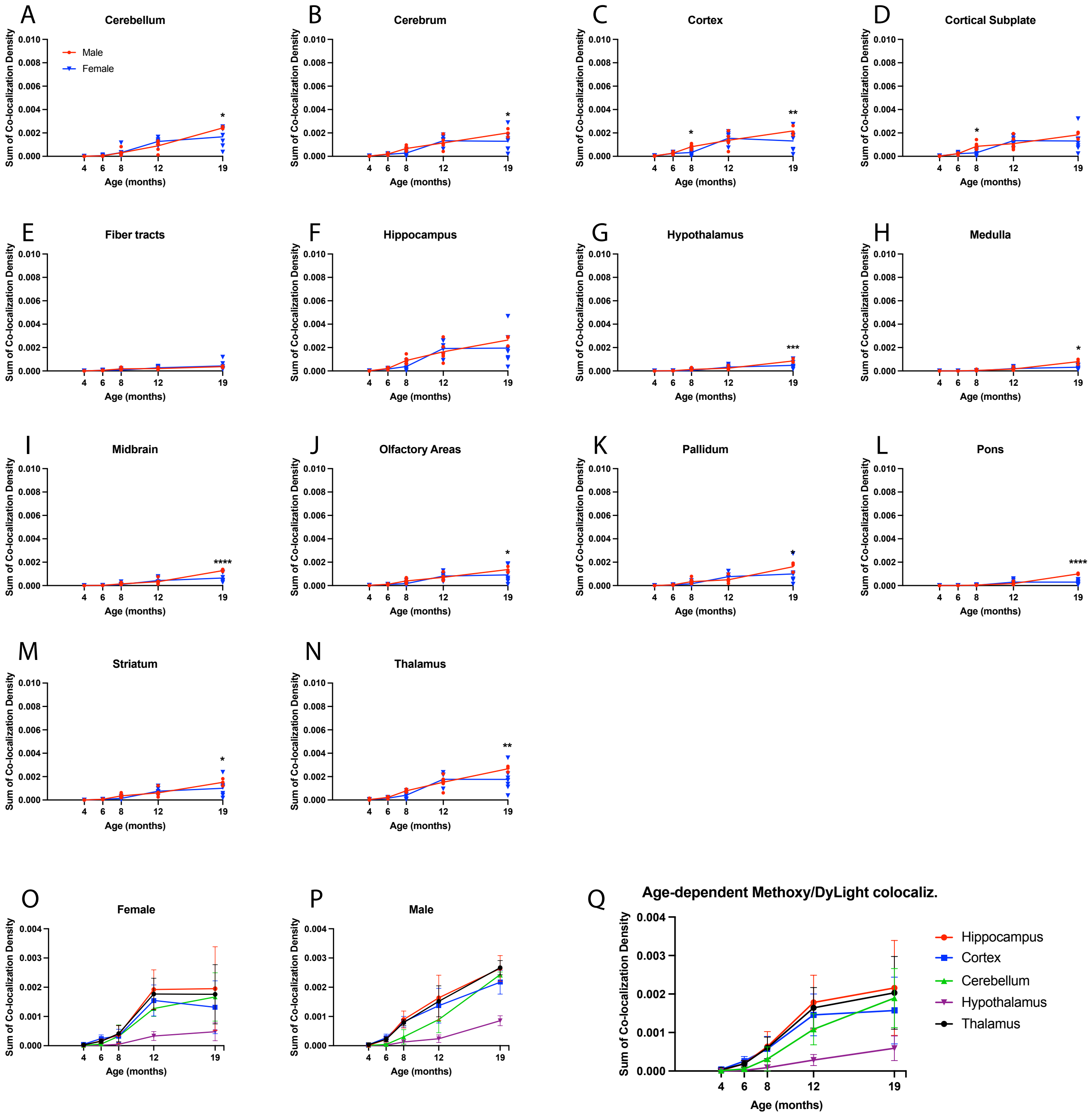


**Supplemental Figure 4. A-N.** Age-dependent, sex-specific analyses of methoxy-X04 co-localization with in vivo DyLight594-stained vasculature in *App*$^{SAA}$ mice in the 13 major brain regions and the cerebrum, according to the Allen Brain Atlas. **O.** Female *App*$^{SAA}$ mice show increased co-localization from 6- to 12-months-of-age before plateauing to 19-months. **P.** Male *App*$^{SAA}$ mice show a linear increase from 6- to 19-months-of-age. **Q.** Combined sexes, age-dependent increase in co-localization density is primarily linear from 4- to 19-months-of-age.

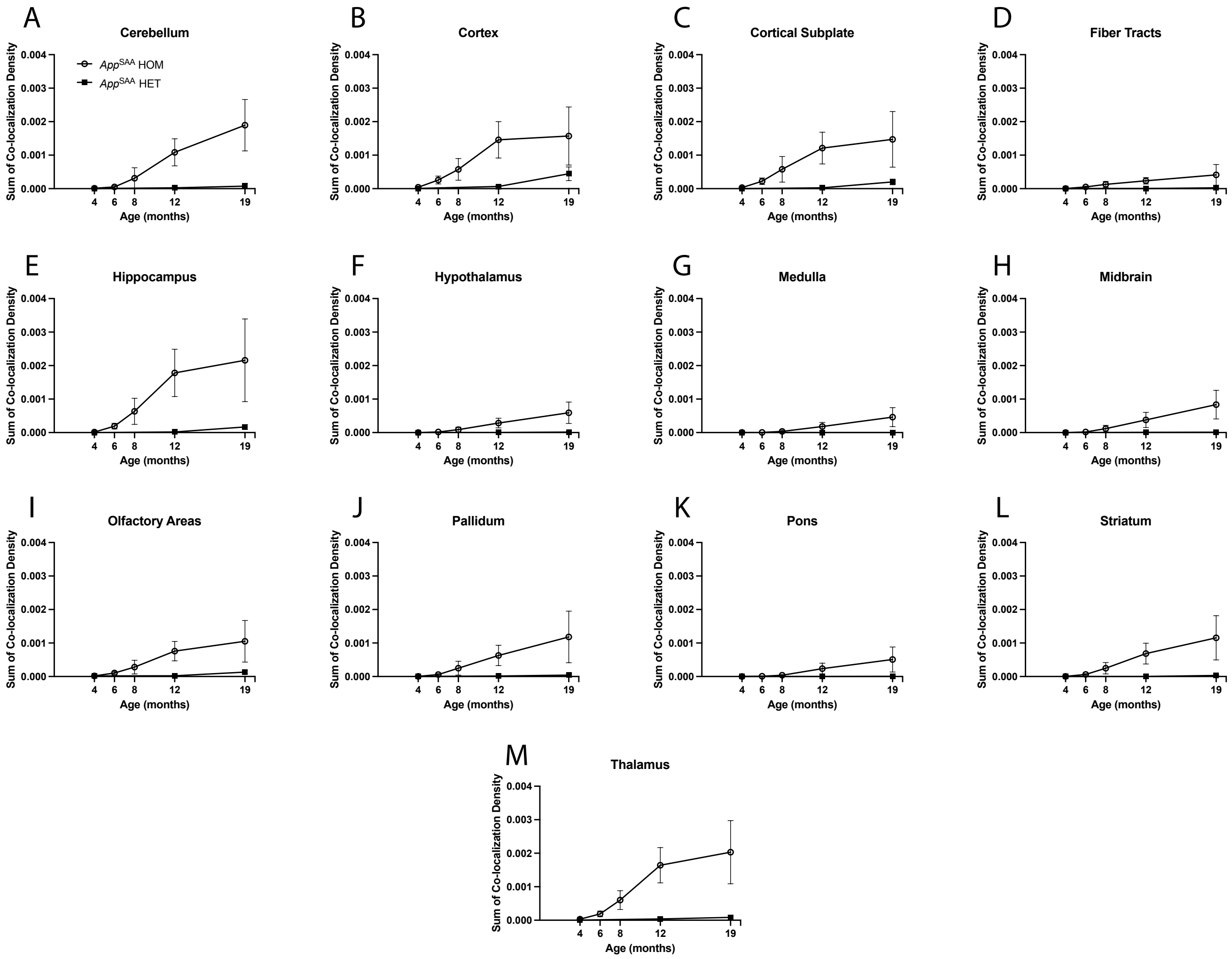


**Supplemental Figure 5. A-N.** Genotype comparisons of co-localization of methoxy-X04 plaques and DyLight594 vasculature between homozygous and heterozygous *App*$^{SAA}$ mice in the 13 major brain regions according to the Allen Brain Atlas. Heterozygous mice show limited co-localization at 19-months-of-age.

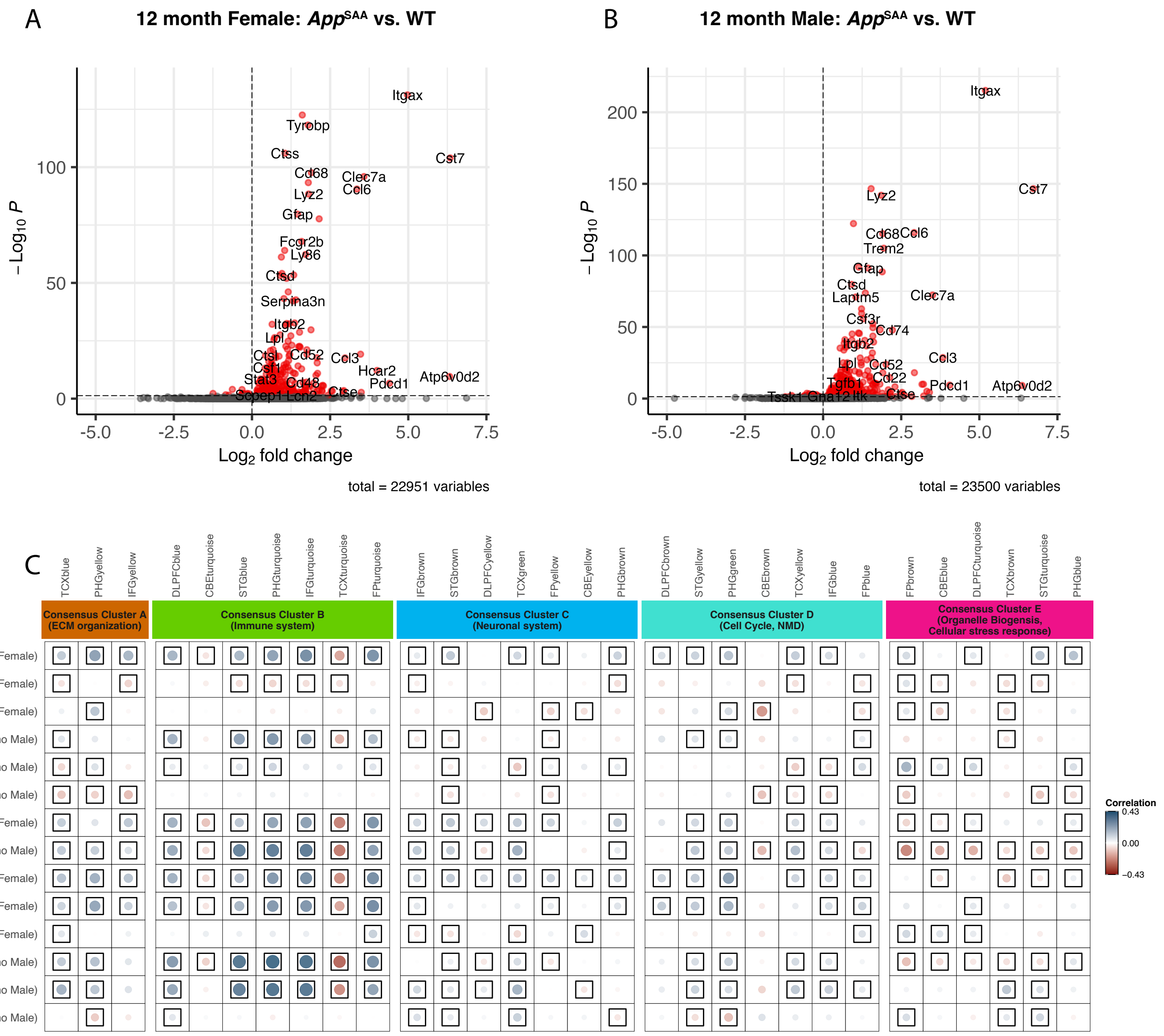


**Supplemental Figure 6.** Additional RNA-seq comparisons. **A-B.** Volcano plot highlighting DEGs from 12-month brain transcriptomic data in **A.** female *App*$^{SAA}$ mice and **B.** male mice compared to B6J controls. **C.** Pearson correlation coefficients for gene expression changes in mice and human disease (log fold change for cases minus controls) comparing male and female 5xFAD mice at 4-, 8-, and 12-months to male and female *App*$^{SAA}$ homozygous and heterozygous mice at 4- and 12-months. The five AMP-AD consensus clusters define the major functional groups of AD-related alterations. Positive correlations are shown in blue and negative correlations in red with color intensity and size of the circles proportional to the correlation coefficient. Boxes indicate statistical significance at p ≤ 0.05.

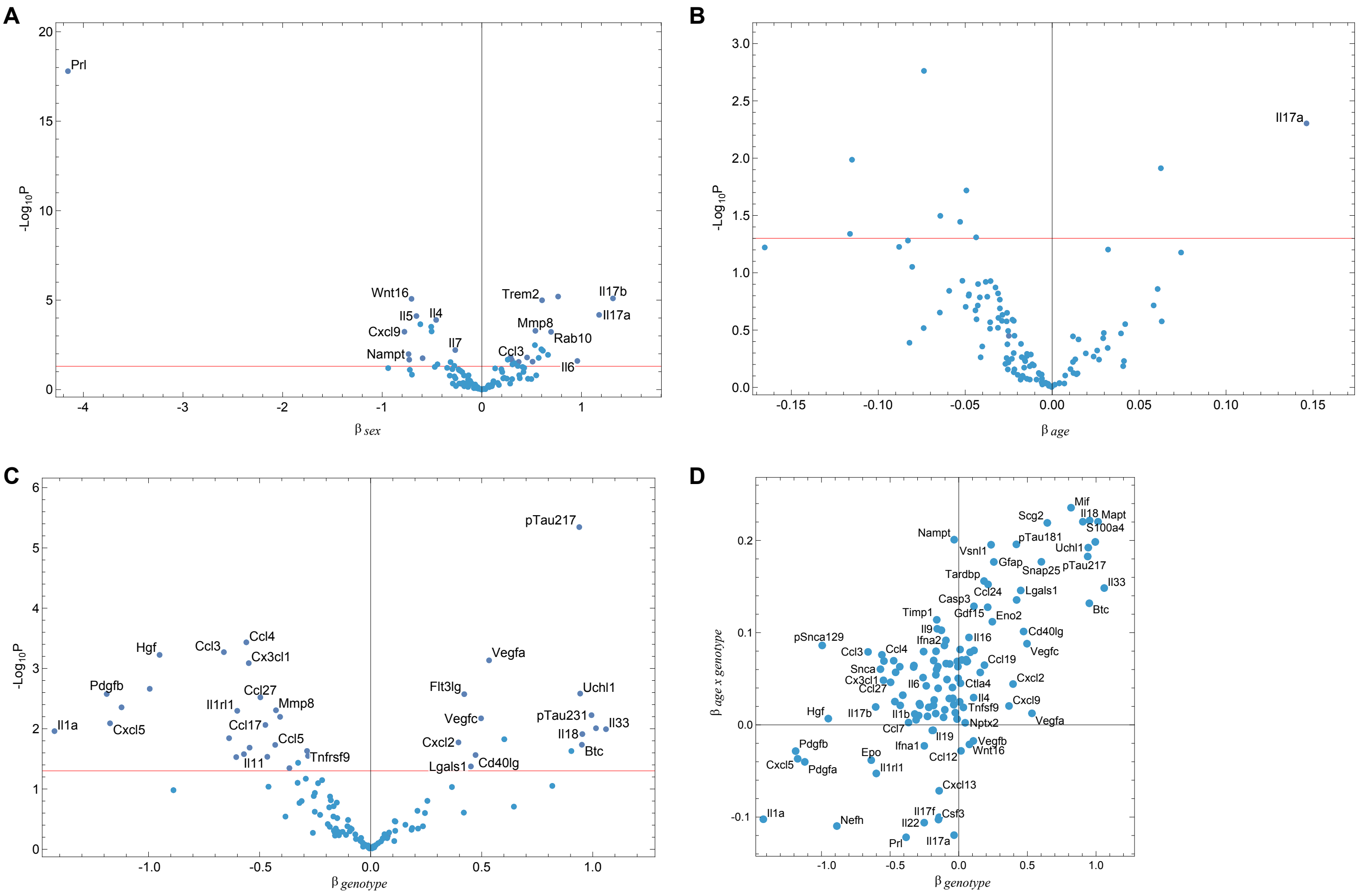


**Supplemental Figure 7.** Effect and p-value estimates from linear modeling of NULISA™ proteins from: **A.** male sex, **B.** age relative to mean animal age, and **C.** *App*$^{SAA}$ genotype. **D.** Correlation of genotype effects and age-by-genotype effects, demonstrating that genotype effects are generally age-dependent.

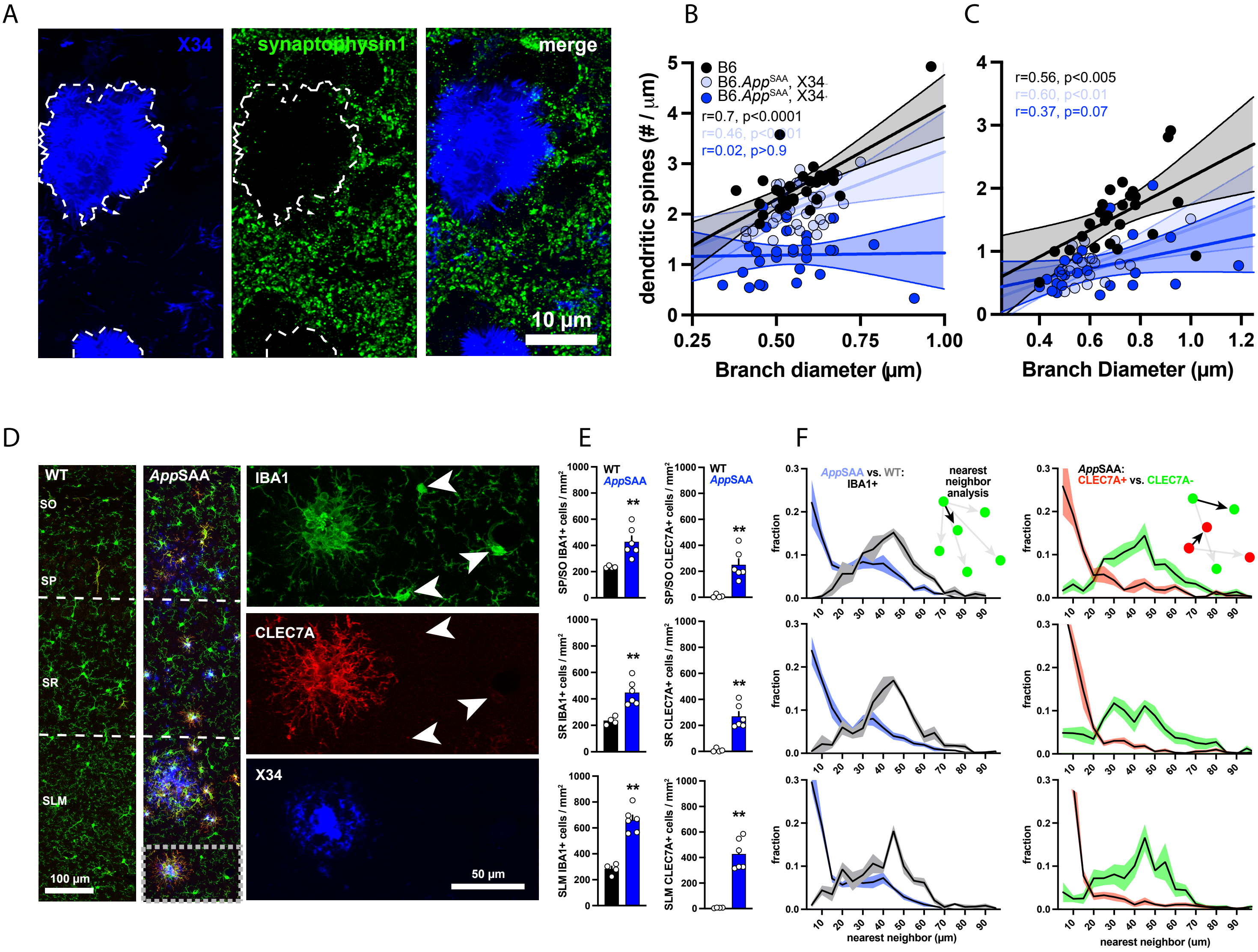


**Supplemental Figure 8.** Synapse loss in 12-month-old *App*$^{SAA}$ mice. **A.** Synaptophysin staining in 12-month-old *App*$^{SAA}$ mice in proximity to X34-labeled plaques is effectively absent. **B.** Correlation plot between branch diameter and spine density on apical oblique dendrites from B6J (black), *App*$^{SAA}$ branches at plaque-free sites (light blue), and *App*$^{SAA}$ branches at X34-plaque+ sites (dark blue). **C.** Same plot for tuft branches. **D.** Immunofluorescent labeling of microglia for IBA-1 and CLEC7A in the in the stratum pyramidale/stratum oriens (SP/SO), stratum radiatum (SR), and stratum moleculare-lacunosum (SLM) of the hippocampus of B6J (WT) and *App*$^{SAA}$ mice. **E.** Nearest neighbor quantification.
